# DNA origami-based single-molecule force spectroscopy unravels the molecular basis of RNA Polymerase III pre-initiation complex stability

**DOI:** 10.1101/775528

**Authors:** Kevin Kramm, Tim Schröder, Jerome Gouge, Andrés Manuel Vera, Florian B. Heiss, Tim Liedl, Christoph Engel, Alessandro Vannini, Philip Tinnefeld, Dina Grohmann

**Author notes:** For correspondence: Dina Grohmann, Department of Biochemistry, Genetics and Microbiology, Institute of Microbiology, University of Regensburg, Universitätsstraße 31, 93053 Regensburg, Germany.

## Abstract

The TATA-binding protein (TBP) and a transcription factor (TF) IIB-like factor compound the fundamental core of all eukaryotic initiation complexes. The reason for the emergence and strict requirement of the additional intiation factor Bdp1, which is unique to the RNA polymerase (RNAP) III sytem, however, remained elusive. A poorly studied aspect in this context is the effect of DNA strain, that arises from DNA compaction and transcriptional activity, on the efficiency of initiation complex formation. We made use of a new nanotechnological tool – a DNA origami-based force clamp - to follow the assembly of human initiation complexes in the Pol II and Pol III system at the single-molecule level under piconewton forces. We demonstrate that TBP-DNA complexes are force-sensitive and TFIIB is necessary and sufficient to stabilise TBP on a strained RNAP II promoter. In contrast, Bdp1 is the pivotal component that ensures stable anchoring of initiation factors, and thus the polymerase itself, in the RNAP III system. Thereby, we offer an explanation for the crucial role of Bdp1 for the high transcriptional output of Pol III genes for the first time.

## Introduction

All cellular life depends on the regulated expression of its genome. The first step in gene expression is transcription, which is carried out by highly conserved multisubunit RNA polymerases (RNAP) that make use of a DNA template to synthesise RNA^1^. Transcription is a cyclic process that can be divided into the initiation, elongation and termination phase. Aided by a number of basal transcription initiation factors, the archaeal-eukaryotic RNAP is recruited to the promoter DNA thereby positioning the RNAP at the transcription start site (TSS) ^2 3^. All archaeal-eukaryotic RNAPs rely on the basal transcription initiation factor TBP and a TFIIB-like factor ^4,5 6,7^, despite some particularities of the Pol I system^8^. TBP is highly conserved in structure and function and recognises an AT-rich DNA stretch, the so-called TATA-box (eukaryotic consensus sequence TATAWAWR with W = T or A and R = G or A ^9^), upstream of the TSS ^10–14^. Canonical binding of TBP to the DNA invokes a 90°C bend in the DNA ^15–17^ when two conserved pairs of phenylalanines are inserted into the promoter DNA between bases 1/2 and 7/8 of the TATA box sequence. Bending leads to a widening of the minor groove of the promoter DNA^16^. TFIIB-like factors associate with the TBP-DNA complex via the C-terminal core domain and concomitantly recognise the B-recognition element (BRE) located adjacent to the TATA-box ^8,18–23^. Even though additional factors (e.g. TFIIE, TFIIH, TFIIF) are involved in the initiation process *in vivo* ^24^, the minimal configuration of TBP and TFIIB factor are sufficient to to recruit the RNAP (in complex with TFIIF) to the promoter in eukaryotic RNAP II transcription system ^25–29^. While the eukaryotic RNAP II system is responsible for the transcription of messenger RNAs and small nucleolar (sn)RNAs, RNAP transcription systems I and III are transcribing ribosomal (r) RNAs and 5S rRNA, U6 snRNA, tRNAs, respectively. The initiation factor setup in the specialised RNAP I and III transcriptions systems, however, diverged from the composition of the RNAP II system and additional initiation factors are required for efficient initiation ^7,8,30^. While TBP was found to be part of the RNAP I initiation machinery *in vivo* ^31–33^, basal transcriptional activity can also be achieved in the absence of TBP ^34–36^ and its functional role in the RNAP I system remains elusive. RNAP III transcription is directed from three different promoter classes that differ in promoter elements and initiation factor requirement ^6,37^. In all cases, transcription initiation in the RNAP III system relies on the multisubunit factor TFIIIB composed of TBP, the TFIIB-like factors Brf1 and Bdp1 (<u>B</u> <u>d</u>ouple <u>p</u>rime or B’’)^7,21^. Bdp1 is unique to RNAP III transcription initiation and has no homologue in the RNAP I or II transcription system. However, Bdp1 is crucially involved in promoter recognition and DNA opening ^38,39^. Vertebrates addionally use a TFIIIB variant that contains Brf2 instead of Brf1. Both factors are structurally similar, but Brf2 binding to the TBP-DNA complex is regulated by the redox state of the cell. The Brf2 containing TFIIIB complex initiates transcription at a small subset of genes, including the selenocysteine tRNA and U6 snRNA. In contrast to Pol II-transcribed snRNA genes, the U6 promoter contains a TATA-box element that is crucial for the specific recruitment of TFIIIB ^38,40,41^. TFIIIB is sufficient for the recruitment of yeast RNAP III *in vitro* ^42^. However, at human type 3 promoters an additional protein complex is involved in transcription initiation, the snRNA activating protein complex (SNAP_c_, reviewed in ^37^).

In addition to biochemical and structural studies, single-molecule fluorescence resonance energy transfer (FRET) and ensemble kinetic studies provided insights into the molecular mechanisms and kinetics of transcription initiation in the archaeal, RNAP II and RNAP III transcription system ^43–52^. Interestingly, TBP-DNA complex lifetimes and bending mechanisms differ significantly between the archaeal and eukaryotic system. Archaeal TBP binds and bends the TATA-DNA only transiently ^44^. In some archaeal systems, TFB is of crucial importance for the recognition of the promoter by TBP ^44^. In all cases, bending is achieved in a single step. Similarly, the interaction of human TBP with the U6 promoter is characterised by short lifetime in the millisecond range^52^ while interaction of yeast TBP with a classical RNAP II promoter is highly stable for minutes to hours and bending occurs in two steps ^44^. TFIIB, e.g., was shown to increase the lifetime of the fully bent state in the RNAP II system. Similarly, the TFIIB-like factor Brf2 prolongs the lifetime of the TBP-DNA complex^52^.

Transcription assays as well as smFRET-based DNA bending assays are performed using naked dsDNA of defined length. *In vivo*, however, transcription initiation factors assemble on the promoter DNA in the context of compact nucleosome structures. As a consequence, the transcriptional landscape in eukaryotes is shaped by chromatin remodelling events ^53^. A number of studies analysed the effect of the nucleosome positioning on transcriptional levels and demonstrated that accesibility of the promoter DNA correlates with transcriptional efficiency ^54^. Another regulative aspect of the nucleosome organisation that has to be considered is the topological effects on DNA introduced by tightly spaced nucleosomes ^55^ and the transcription (and replication) machinery. In this context, DNA is subject to mechanical forces. The effect of these forces on transcription initiation, however, has not been analysed as suitable methodological tools were not available so far. Standard force-sensitive methods like magnetic and optical tweezers require long DNA linker strands that connect the DNA under investigation to the macromolecular world, e.g. in magnetical or optical tweezer experimentes a topological change of the investigated DNA can only be transmitted to the beads via this linker. This in turn contributes to a considerable noise in a tweezer experiment. Consequently, subtle changes in DNA topology introduced by DNA-binding proteins like TBP are extremely difficult to detect ^56^.

Here, we utilise a recently developed DNA origami-based force clamp ^57^ to monitor the influence of DNA strain on the assembly of transcription initiation factors from the human RNAP II and RNAP III transcription system on the promoter DNA. Our data establishes the RNAPIII - specific initiation factor Bdp1 as the pivotal component of the RNAP III initiation complex that ensures stable anchoring of the initiation factor TFIIIB, and by extension the RNAP III, at the promoter. This exceptional stability provides a stable anchor point for RNAP III at the promoter that’s supports the transcription of the short U6, tRNA and 5S rRNAs. Moreover, we demonstrate for the first time that the DNA origami force clamp is a powerful tool to study the force-dependency of complex protein assemblies and that this nanoscopic tool provides detailed mechanistic and kinetic information about biological processes that have not been accessible before.

## Results

### DNA origami-based force clamp to probe force sensitivity of transcription initiation complexes

Recently, we introduced a DNA origami-based force clamp that exerts forces in the piconewton regime on a DNA segment (**Figure 1A**)^57^. This nanosized force clamp exploits the entropic spring behaviour of single-stranded DNA (ssDNA) that is placed in the middle of the DNA origami clamp. Forces are tunable by adjusting the length of the ssDNA that is connected to the rigid body of the DNA origami thereby providing two fixed anchor points for the ssDNA (**Figure 1B**). Due to the reduced conformational freedom of a short DNA segment (equivalent with a reduced entropy of the system), higher strain (e.g. force) acts on the DNA. The resulting forces were calculated using a modified freely jointed chain model^57,58^ (for details see **Supplementary Methods**). In this study, we employed DNA origami force clamps with forces ranging from 0 to 6.6 pN. The major advantage of the nanoscopic force clamp is that it acts autonomously and does not require a physical connection to a macroscopic instrument. Moreover, the DNA origami force clamp can be produced and used in a highly parallelised manner. In order to study the force-dependency of transcription initiation factor assembly on the promoter DNA, we engineered a prototypical RNAPII (Adenovirus major late promoter, AdMLP) and RNAPIII promoter (human U6 snRNA promoter) sequence into the DNA origami (**Supplementary Figure 1**). The AdMLP promoter contains a TATA-box and BRE element sequence, which are targeted by TBP and TFIIB, respectively. The TATA-box of the U6 snRNA promoter is flanked by the GR-element at position −3/−4 and TD-motif at position +3/+4 relative to the TATA-box (**Supplementary Figure 1**), which are bound by the TFIIB-like factor Brf2 ^52^. Annealing of a short complementary additional DNA strand that carries a donor (Atto532) and acceptor (Atto647N) fluorophore allows the detection of TBP-induced DNA binding via smFRET measurements (**Figure 1C** and **Supplementary Figure 1**). The correct folding of the DNA nanostructure was verified using transmission electron microscopy (**Supplementary Figure 2**). The successful hybridisation of the fluorescently labelled DNA strand is demonstrated by fluorescence correlation spectroscopy measurements as the short dsDNA promoter diffuses seven times faster than the respective DNA origami where the labelled DNA is part of the high molecular weight DNA origami structure (Supplementary Figure 3). We first performed smFRET measurements on freely diffusing DNA origamis and found a single uniform low FRET population for all forces for the AdMLP and U6 promoter force clamps (**Figure 2 and 3**). The measured FRET efficiencies are in good agreement with FRET efficiencies obtained from linear dsDNA promoter DNAs (**Supplementary Figure 4**). This demonstrates that the conformation of the promoter DNA is not significantly changed when it is incorporated into the DNA origami force clamp and forces are applied.

**Figure 1:**
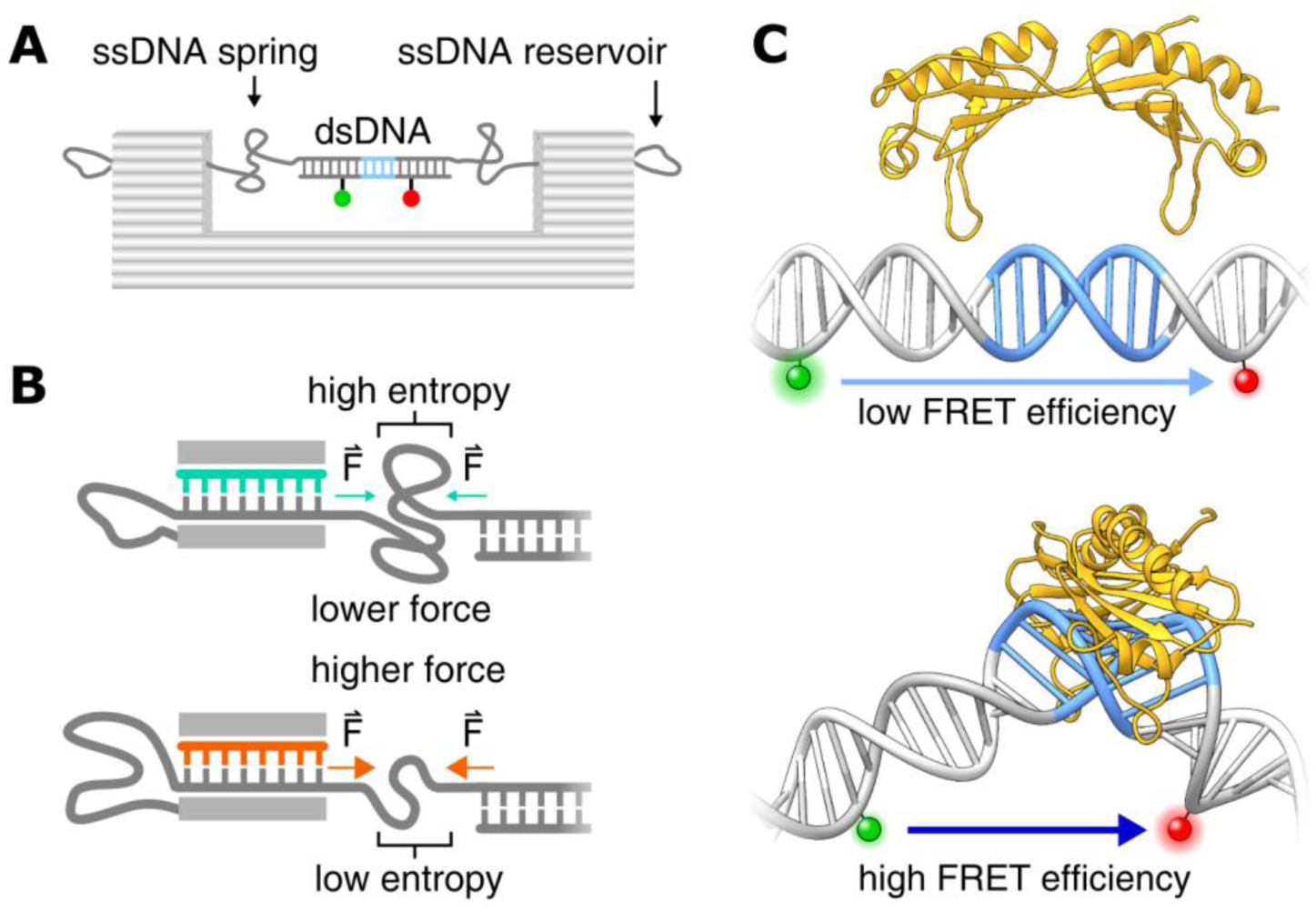
DNA-origami based force clamp monitors TBP-induced DNA bending under force. **A)** Schematic overview of the DNA origami force clamp. The ssDNA spring protrudes from the DNA origami body and spans the 43 nm gap of the rigid DNA origami clamp body (grey). Centered withing the ssDNA spring is a double stranded promoter region incorporating the TATA-box element (blue) flanked by a donor/acceptor (green/red) fluorescent dye pair for FRET sensing. **B)** The ssDNA spring length can be adjusted with DNA from the reservoir by using different staples (teal/orange) during assembly. Reducing the number of nucleotides spanning the gap leads to a smaller number of adoptable conformations of the ssDNA chain and thus results in a higher entropic force. **C)** Single-pair FRET assay as readout for the bending of promoter DNA by the TATA-binding protein (TBP, yellow). A donor (ATTO 532, green) and acceptor fluorophore (ATTO 647N, red) flank the TATA-box element (blue) resulting in a low efficiency Förster resonance energy transfer (FRET) between both dyes. Binding of TBP bends the DNA by approximately 90° thereby decreasing the distance between the fluorophors resulting in an increase in FRET efficiency (DNA-TBP structures adapted from: PDB 5FUR).

**Figure 2:**
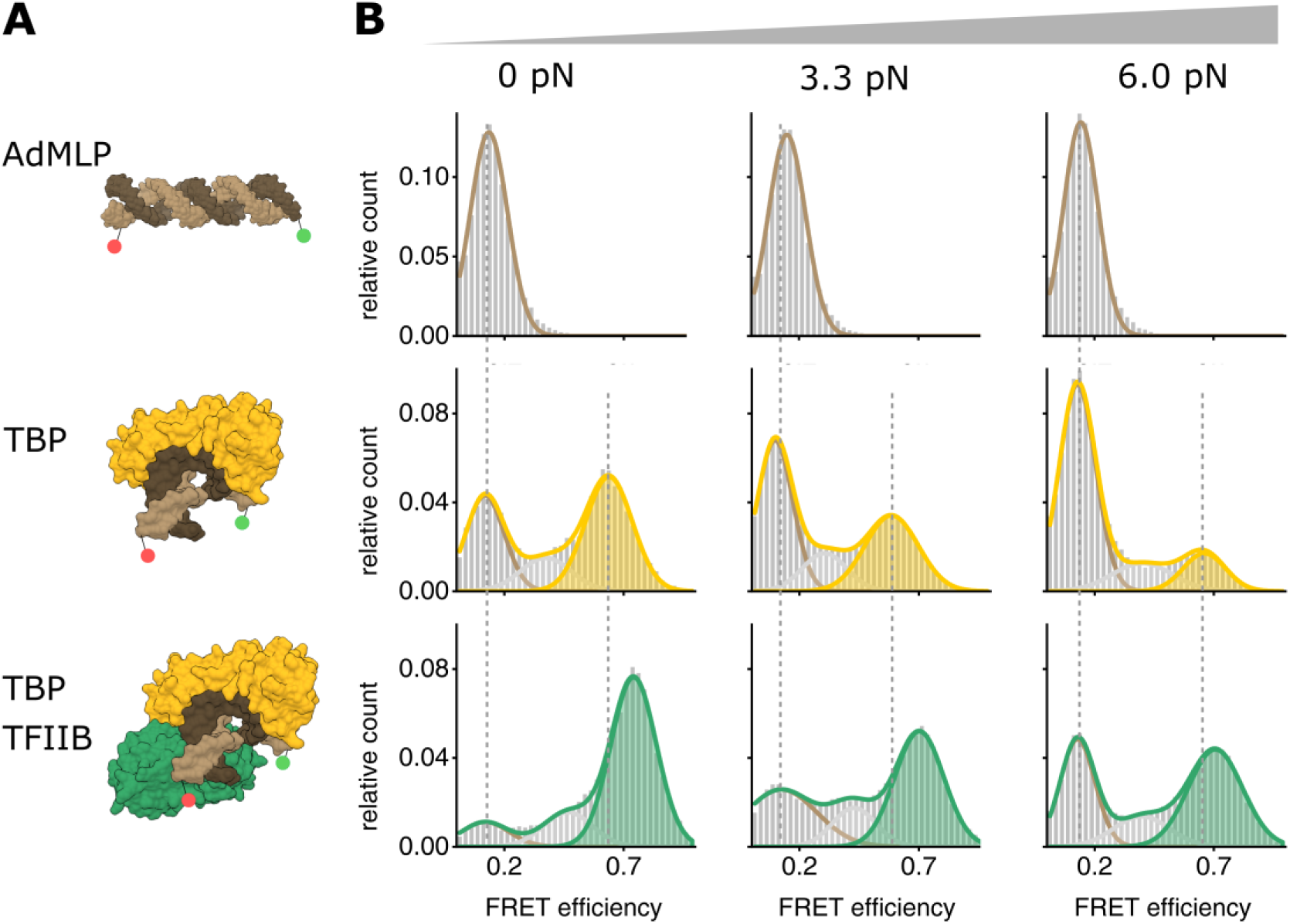
Force dependency of promoter binding of RNA polymerase II initiation factors at a canonical Pol II promoter. **A)** Structural model (PDB: 1C9B) of the adenovirus major late promoter (AdMLP, brown) in an unbent conformation and the 90° bend state bound by TBP (yellow) and TFIIB (green). **B)** Single-molecule FRET measurements monitor TBP-induced DNA bending after addition of TBP (20 nM) or TBP and TFIIB (200 nM) to the AdMLP DNA origami force clamps at increasing forces (0, 3.3, 6.0 pN). FRET efficiency histograms showing the relative distribution between the unbent DNA state (low FRET state, E = 0.12, brown) and TBP-induced bent state (high FRET population, E = 0..63 (TBP only, yellow), E = 0.72 (TBP/TFIIB, green)). Low and high FRET populations were fitted with a Gaussian distribution. Each measurement was carried out at least three times. See also **Supplementary Figure 5** and **Supplmenetary Table 4**.

**Figure 3:**
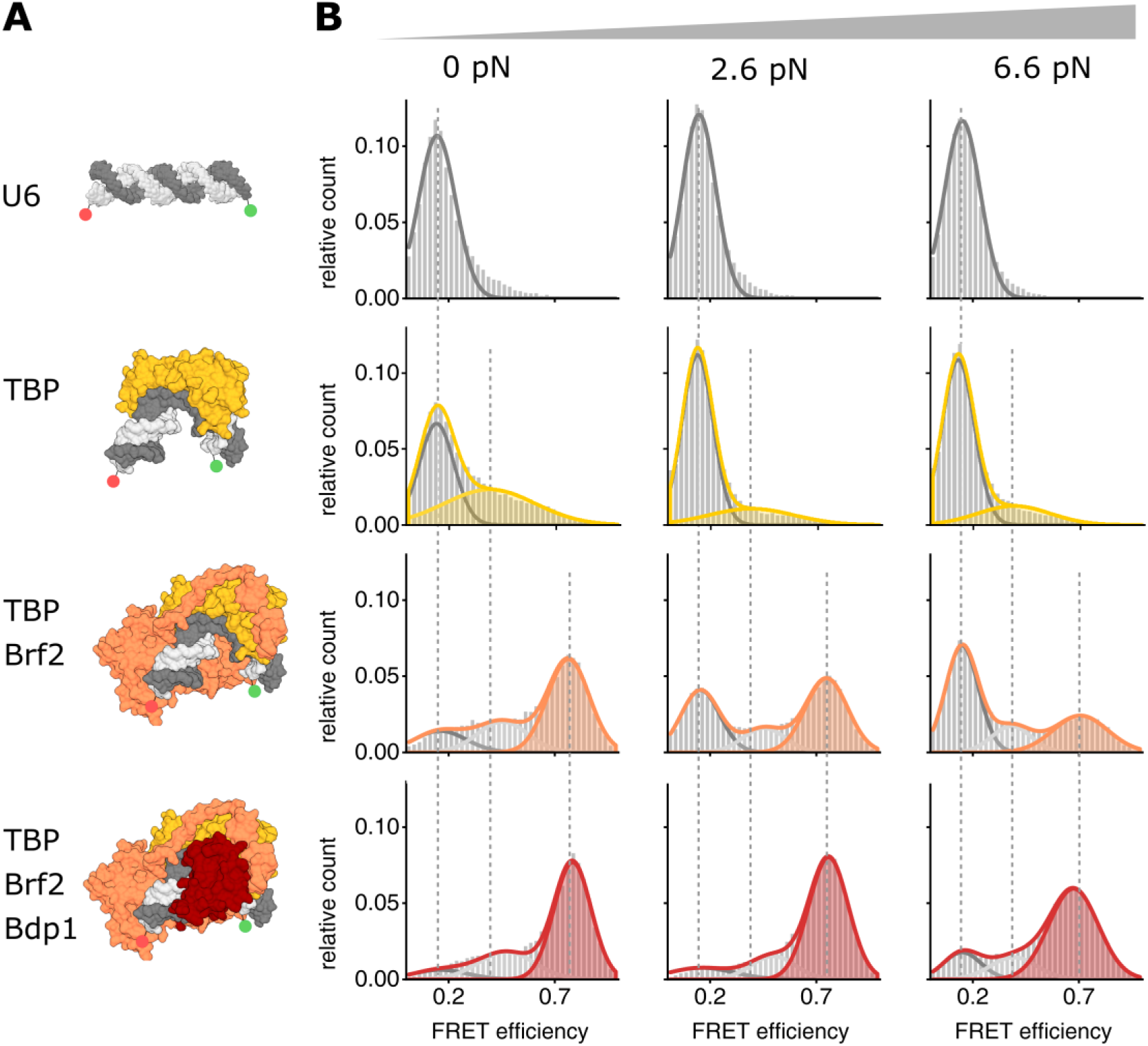
Force dependency of promoter binding of RNA polymerase III initiation factors at a canonical Pol III promoter. **A)** Structural model (PDB: 5N9G) of the U6 snRNA promoter (U6, dark grey) in an unbent conformation and the 90° bent state bound by TBP (yellow), TBP+Brf2 (orange) and TBP+Brf2+Bdp1 (red). **B)** Single-molecule FRET measurements monitor TBP-induced DNA bending after addition of TBP (20 nM), TBP/Brf2 (20 nM) or TBP/Brf2/Bdp1 (20 nM) to the AdMLP DNA origami force clamps at increasing forces (0, 2.6, 6.6 pN). FRET efficiency histograms showing the relative distribution between the unbent DNA state (low FRET state, E = 0.19, grey) and TBP-induced bent states in the absence and presence of additional initiation factors (high FRET population, E = 0.39 (TBP only, yellow), E = 0.75 (TBP/Brf2, orange), E = 0.76 (TBP/Brf2/Bdp1, red)). Low and high FRET populations were fitted with a Gaussian distribution. Each measurement was carried out at least three times. See also **Supplementary Figure 5** and **Supplementary Table 4**.

### TBP-induced promoter DNA bending of a Pol II and Pol III promoter under force

First, we probed the force-dependency of the human RNAP II transcription initiation complex formation. Basal transcription levels in the RNAPII transcription initiation can be achieved using TBP and TFIIB only. Hence, we added TBP or TBP/TFIIB to the DNA origami force clamp that carries a canonical RNAPII promoter (AdMLP promoter). At the TBP concentration chosen (20 nM), 50% of the molecules showed a high FRET value with a FRET efficiency of 0.63 at 0 pN (**Figure 2, Supplementary Figure 5** and **Supplementary Table 3**). Similar results were obtained using linear dsDNA demonstrating that the DNA origami force clamp is suited to probe TBP-induced DNA bending (**Supplementary Figure 4**). An increase in force to 3.3 and 6.0 pN resulted in a decrease in the fraction of the high FRET population with only 15% of the molecules in the high FRET state at 6.0 pN (**Figure 2, Supplementary Figure 5** and **Supplementary Table 3**). These data show that the bending of a RNAP II promoter by TBP is force-dependent. Similarly, addition of human TBP to the U6 promoter DNA origami led to the appearance of a high FRET population (E = 0.39) while the fraction of the U6 promoter by TBP is reduced at higher forces (**Figure 3**; this data will be discussed in detail below).

### TFIIB and Brf2/Bdp1 are required to establish fully stable Pol II and Pol III initiation complexes

Addition of TFIIB changes the equilibirium between the bent and unbent DNA state dramatically. At 0 pN almost all molecules were found in the high FRET state (Figure 2). Increasing the force to 3.3 and 6.0 pN resulted in a decreased high FRET population. However, at 6.0 pN a significantly higher fraction of molecules (49%) exhibited a high FRET state as compared to the samples that only contained TBP. Moreover, the high FRET is shifted to a value of E = 0.72 indicating that the bending angle is slightly increased in the presence of TFIIB. These results suggest that TFIIB significantly stabilises the TBP-DNA interaction, which is in agreement with previous smFRET studies that showed that TFIIB not only extends the TBP-DNA complex lifetimes but also shifts the equilibrium towards the fully bent state^44^.

Addition of the TFIIB-like factor Brf2 to the TBP-U6 promoter complex also resulted in a stabilisation of the TBP-DNA complex and a shift of the bent DNA population to a higher FRET efficiency (E = 0.74). In both cases, however, the complex was still force-sensitive and only a small fraction (Brf2 31%, TFIIB 30%) of molecules was found in the bent state at 6.6 pN (**Figure 2, Supplementary Figure 5**). In previous studies, we observed a significant increase in the lifetime of the complexes when Brf2 was added to the TBP-DNA complex^52^. Addition of Bdp1 to the TBP-Brf2-DNA complex, however, did not substantially affect the complex lifetime when linear promoter DNA was used for smFRET measurements^52^. Hence, we wondered whether Bdp1 influences the complex stability when the DNA experienced increased strain. Probing the force-sensitivity of the TBP-Brf2-Bdp1-DNA complex showed that even at 6.6 pN, the majority of molecules (69%) was found in a bent DNA state. We therefore conclude that in the Pol III system, Bdp1 is the decisive initiation factor that renders the initiation complex fully stable (**Figure 3**). In contrast, TFIIB suffices to ensure such a stable complex formation in the Pol II system.

### Increased DNA strain destabilises the TBP-DNA interaction

Previous measurements showed that the TBP-DNA interaction is dynamic ^44,52^. This gave us the opportunity to ask whether the increase in strain reduces the lifetime of the TBP-DNA complex (enhanced TBP dissociation with increase in force) or prolonges the lifetime of the unbent DNA state (inhibited TBP association with increase in force). To answer this question, we use two different strategies adapted to the underlying kinetics of association/dissociation process. Slow kinetics in the minutes time regime were measured by acquiring smFRET distributions at different time points after mixing the constituents of the transcription complex. Faster kinetics were measured by monitoring the high-FRET and low FRET state lifetimes directly on single immobilized complexes. Time-resolved smFRET measurements of the TBP-DNA interaction at 0 and 6.0 (AdMLP) or 6.6 pN (U6 promoter) showed that the TBP-AdMLP promoter exhibits a lifetime of 311 ± 62 s at 0 pN force while the interaction between TBP and the U6 promoter is short-lived (τ_bent_ = 0.54 ± 0.02 s) (**Figure 4** and **Table S3**). This is in agreement with previous observations using linear dsDNAs ^52^. Increased force leads to an increase in the lifetime of the unbent state while the lifetime of the bent state remains constant (AdMLP: τ_unbent_ = 844 ± 149 s and τ_bent_ = 312 ± 52 s). Higher forces do not influence the lifetime of the unbent state in case of the TBP-U6 promoter DNA complex (0 pN: τ_unbent_ = 0.21 ± 0.01 s, 6.6 pN: τ_unbent_ = 0.26 ± 0.01 s). However, the lifetime of the bent state is slightly reduced at 6.6 pN as compared to 0 pN (0 pN: τ_bent_ = 0.54 ± 0.02 s and τ_bent_ = 0.35 ± 0.01 s). These data suggest that two factors contribute to the reduction of the bent DNA states at higher forces: i) destabilisation of the TBP-DNA state with increased propability of TBP dissociation from the DNA at higher forces (spring-loaded TBP ejection mechanism) in case of the U6 promoter and ii) a decreased propability of the TBP to form a stable complex with DNA (TBP entry denial) at the AdML promoter. It seems plausible that DNA under strain does provide less flexibility between the bases for the two phenylalanines pairs to insert into the DNA and thereby entry of TBP into the DNA is denied. The interaction of the already inserted phenylalanines with the bases of the DNA, on the other hand, may be reduced at higher DNA strain, leading to ejection at high strains.

**Figure 4:**
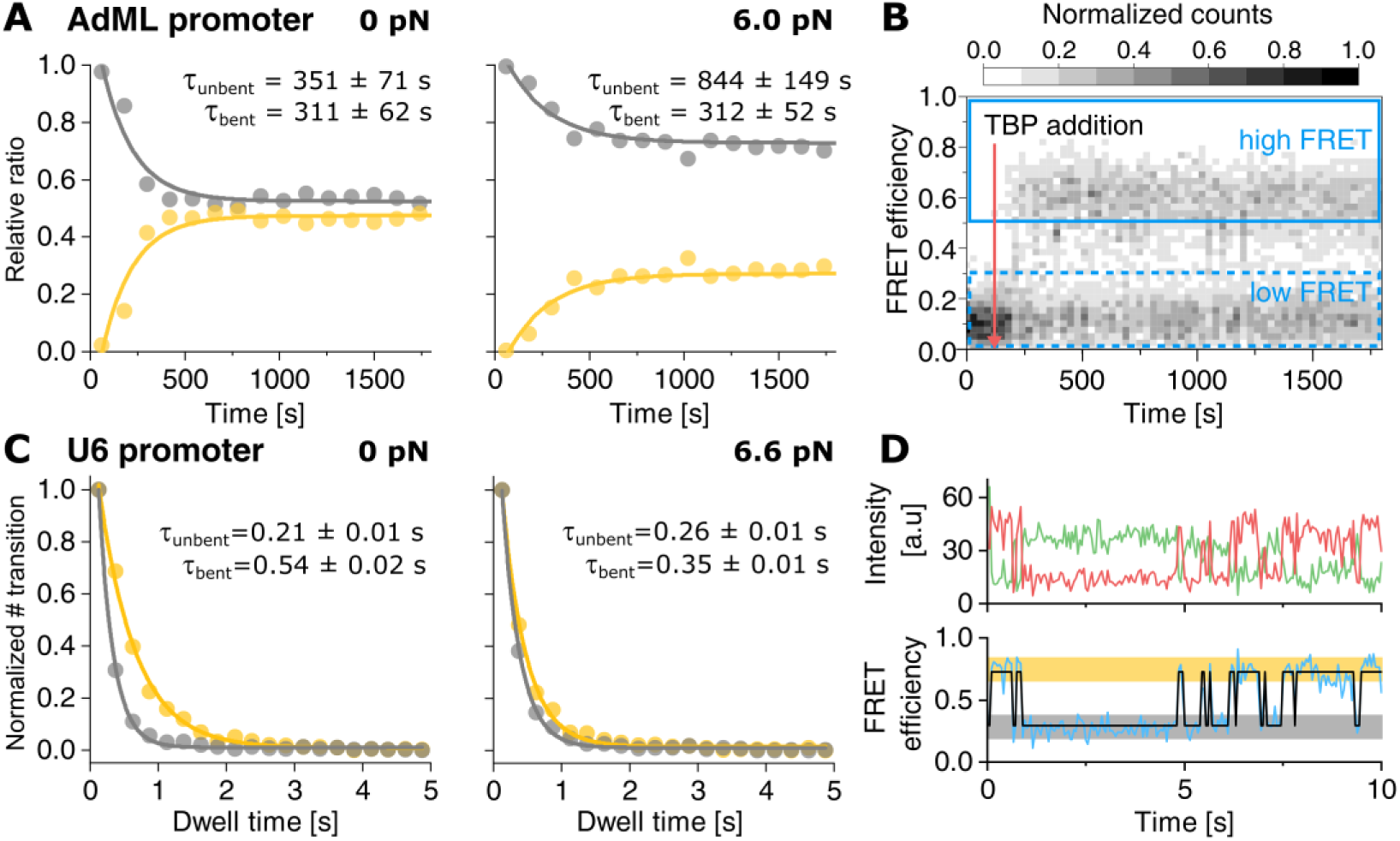
Kinetic analysis of the influence of force on TBP-induced DNA-bending. **A)** Relative ratios of low FRET (unbound DNA, grey) to high FRET state (TBP-DNA complex, yellow) in a kinetic experiment showing the relative fraction of the TBP-DNA complex to the unbent U6 promoter at 0 pN and 6.0 pN. Dwell times were calculated by deconvolution with a perturbation-relaxation model. Data were fitted with a mono-exponential function. **B)** Representative FRET efficiency-time plot of a time course experiment at 0 pN force. TBP (20 nM) was added at 2 min (red arrow). Areas used for calculating the ratio of low and high FRET are indicated by blue boxes. **C)** Dwell-time histograms of the U6 promoter in the unbent (grey) and bent (yellow) state at 0 pN and 6.6 pN force. **D**) Representative donor (green) and acceptor (red) intensity-time trace and the resulting FRET efficiency (blue) fitted with the idealised two-state trace (black) of TBP binding to the U6 promoter at 0 pN force. The low FRET and high FRET states are highlighted in grey and yellow, respectively. Values are given as mean ± s.e.m. (see also **Supplementary Figure 6** and **Table S3**).

## Discussion

During the initiation phase of transcription, the transcriptional machinery is assembled at the promoter. The minimal factor requirement for transcription initiation consists of TBP and TFIIB to recruit RNAP II and TBP, Brf1 or Brf2 and Bdp1 and additionally SNAPc to recruit RNAP III. One of the interesting questions in this context is why the RNAP III machinery relies on a third basal initiation factor not conserved in the RNAP I or RNAP II system? Based on our data, part of the answer might be found in the fact that promoter DNA - rather than being a rigid stick-like molecule - is part of a complex chromatin superstructure with dynamic structural variability and consequently subject to mechanical forces in the dynamic landscape of chromatin that is constantly exposed to changes by chromatin remodelers and gene activators ^54^. This also includes loop formation and tight nucleosomal packaging that exerts mechanical forces on the DNA ^59,60^. Additionally, attractive interaction between nucleosomes mediated by the histone tail domains have recently been observed using DNA nanotechnology ^55^. These close-range interactions vary in strength between −0.3 to −8 kcal/mol which falls into the range covered by our experiments ^55,61–64^ (**Supplementary Figure 7**). However, the chromatin landscape and consequently the forces that act on the promoter DNA differ between Pol II and III promoters. In this work, we investigated the force sensitivity of transcription initiation factor assembly at the promoter DNA at variable forces employing a novel method to carry out force measurements based on a DNA origami force clamp^57^. Combined with a smFRET assay, we were able to quantify TBP-induced promoter DNA bending and to evaluate the influence of additional initiation factors. Using identical TBP concentrations, we found that human TBP bends the U6 snRNA promoter less efficiently under force than the AdML promoter. This is not surprising as only four out of eight bases of the TATA sequence of the U6 promoter sequence match the human consensus TATA box sequence^9^. In contrast, the AdMLP provides a perfect TATA box. This is also reflected in the bent/unbent state lifetime measured for both complexes (Figure 4). Here, mainly the unbent state lifetime increases with force, thus the AdMLP-TBP complex with its higher lifetime is less effected than the transient U6-TBP complex. Our data show that TBP in conjunction with TFIIB forms stable and force-resistant complexes at the prototypical RNAP II AdML promoter. The long lifetime of the TBP-DNA complex, the observed stabilising effect of TFIIB and the increase in bending angle upon addition of TFIIB is consistent with previous smFRET measurements using yeast TBP/TFIIB ^44^. In the RNAP III transcription system, we observed that the TFIIB-like factor Brf2 also enhances the stability of the TBP-DNA complex ^52^. Interestingly, the addition of the third initiation factor, Bdp1, yields an outstandingly stable initiation complex at the U6 promoter. It is noteworthy that the spliceosomal U6 RNA and other RNAP III gene products are highly expressed. This in turn requires robust formation of initiation complexes at the promoter as transcriptional regulation cannot take place at the level of elongation at these extremely short genes. Hence, the RNAP III-exclusive initiation factor Bdp1 plays a decisive role in transcription initiation as it allows the maintenance of fully assembled TFIIIB - promoter DNA complex. The stable anchoring of initiation proteins as well as the RNAP III is furthermore of crucial importance as RNAP III is thought to undergo extensive cycles of facilitated re-initiation ^65–67^. RNAP III only transcribes very short RNAs (5S rRNA, tRNAs, U6 snRNA) and biochemical and recent structural data suggest that RNAP III, in contrast to RNAP II, might not escape from the promoter during transcription elongation but possibly remains bound to the promoter and re-initiates directly after termination ^65,67 66,68^. Hence, initiation factors at the promoter are situated at a DNA section that is topological restrained on the one hand side by the −1 nucleosome, which is stably positioned at −150 bp ^69^ and a firmly associated transcribing RNA polymerase. Upon promoter opening of the DNA by RNAP III in concert with Bdp1, the DNA section experiences torsional strain as the DNA is unwound and the strain cannot be released due to the static nucleosome and RNAP III that represent fixed boundaries (**Figure 5**). Hence, TFIIIB is likely to experience mechanical forces that are compensated by the extremely stable initiation complex. Moreover, in a model where the polymerase remains bound to the promoter, strain would build up during transcription between the promoter binding site and the active site due to the increasing amount of transcribed DNA that has to be accommodated in the polymerase. This addtionally increases the forces that the transcription initiation complex has to withstand.

**Figure 5:**
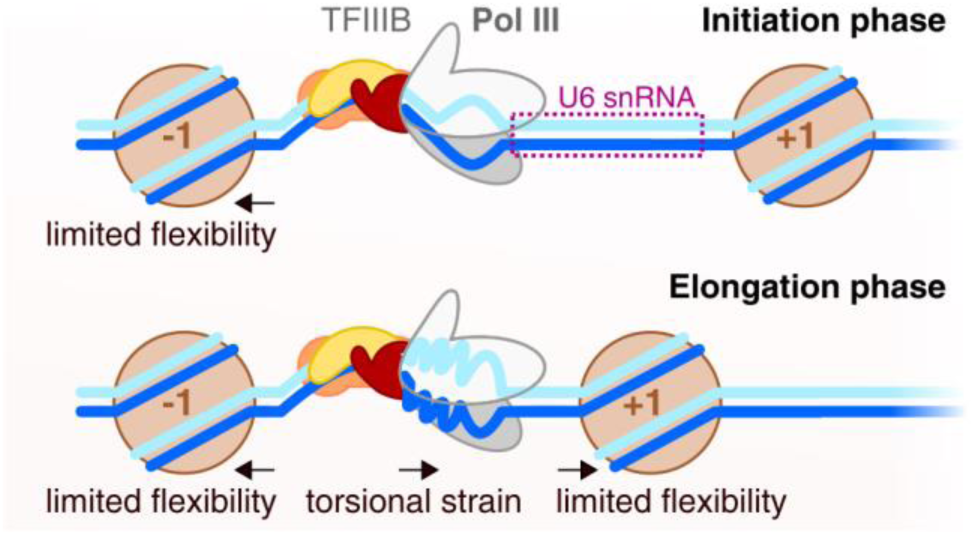
Model describing the role of DNA strain in RNA polymerase III initiation complexes. On the U6 snRNA promoter, the −1 nucleosome is firmly positioned close to the upstream promoter region, limiting DNA flexibility. Continuous transcription of the U6 snRNA by promoter-bound RNA polymerase III creates torsional strain. The +1 nucleosome is positioned downstream of the gene body. Proteins and nucleic acids are color coded as follows: TBP (yellow), TFIIB (green), Brf2 (orange), Bdp1 (red), RNA polymerase II/III (grey), nucleosome (brown), template strand (blue), non-template strand (cyan).

The situation is different at RNAP II promoters as RNAP II transcribes mRNAs that can be hundreds of basepairs in length and re-initiation does not seem to play a role. Another point to consider is that RNAP II and III promoters display a nucleosome depleted region around the transcription start site (TSS) but a conserved +1 nucleosome is found at position +40 in genes with elongating RNAP and +10 in silent genes (RNAP II) ^70,71^ and 220 bp (RNAP III)^69^. As the position of the +1 nucleosome does not show a strong sequence-dependency and its position appears to be flexible when nucleosomes are reconstituted on naked DNA *in vitro* ^*71*^, it has been speculated that initiation factors situated at the promoter help to establish the position of the +1 nucleosome ^54,72^. This might be especially relevant for RNAP II genes where the +1 nucleosome is found in close proximity to the TSS. In this case, initiation factors need to be stably attached at the promoter in order to avoid displacement by the nucleosome. Here, TFIIB acts as the initial stabilising factor at RNAP II promoters to secure TBP at the DNA and this minimal initiation complex can be further extended by additional initiation factors like TFIIA and ultimately extended to include the Mediator complex^19^. Homologues factors are not found in the RNAP III system but our studies show that the addition of Bdp1 to the RNAP III initiation factor lineup is necessary and sufficient to maintain an active initiation complex even when the transcribing RNAP III potentially causes increased DNA strain in the promoter DNA. Interestingly, extending the initial TBP-DNA complex by aditional transcription factors increases the lifetime of the unbent state increases with force. This indicates that the tension on the DNA is a mechanism of gene regulation. The packaging, histone placement, action of the replication machinery and binding of regulatory proteins will certainly have an impact on the tension that the iniation complex is exposed to. Thus, besides steric effects, tension influences transcription. On the other hand, after the transcription initiation complex has formed (i.e. more than one transcription factor is assembled at the promoter), the lifetime of the complex becomes independent of force. This might indicate that after the decision for transcription was taken, the process should become independent of mechanical factors ensuring that the RNA polymerase enters the elongation phase of transcription.

## Material and Methods

### Proteins

All proteins were expressed and purified as described previously ^52^. For the measurements shown we used a N-terminal Bdp1 variant that encompasses residues 130-484 that efficiently forms a complex composed of TBP, Brf2 and promoter DNA^21,52^.

### Cloning of promoter DNA sequences into the M13 DNA origami scaffold

The Force-clamp origami used in this work is based on the M13mp18 ssDNA. The multicloning site of the ssDNA phage DNA is located within the spring region of the force clamp, and the two different RNA polymerase promoters were cloned between the BamHI-HindIII restriction sites of the multicloning site. AdMPL promoter and U6sn RNA promoters were assembled by means of hybridisation of 5’-phosphorylated forward and reverse oligonucleotides (**Supplementary Table 1**). Annealing of the forward and reverse oligonucleotides generate BamHI and HindIII sticky ends. Cloning was performed using the replicative form (dsDNA) of the M13 phage, and high titer phage stocks and ssDNA M13 DNA for DNA origami assembly were prepared as described elsewhere ^73,74^. In both cases, promoter sequences were confirmed by sequencing after cloning.

### Preparation of doubly labelled single-stranded DNAs

Doubly labelled single-stranded DNAs were prepared from individual DNA strands that carry either the donor or the acceptor fluorophore (**Supplementary Table 1**). The final DNA strand carries both dyes and is complementary to the promoter region of the origami scaffold. 10 µM of the appropriate donor strand (_D), acceptor strand (_A) and complementary ligation strand (_Lig) were hybridised in 100 µL annealing buffer (Tris HCl pH 8.0, 150 mM NaCl), heated to 90 °C for 3 min and cooled down to 20 °C over 2 h. For the ligation, 20 µL 10x T4 ligase buffer (NEB), 70 µL Millipore water and 10 µL T4 DNA ligase (NEB) were added to the hybridization reaction and incubated for 60 min at 20 °C.

In order to purifiy the ligated single strand DNA, the DNA was separated on a 200 µL preparative denaturing TBE gel (15% (v/v) acrylamide/bisacrylamid (19:1), 6 M urea). To this end, RNA loading buffer (47.5 % glycerol (v/v) 0.1 % (v/v) SDS, 0.5 mM EDTA) was added to the ligation reaction and the sample was heated to 80°C and cooled on ice. The DNA was separated at 200 V over 40 min. The gel was visualized under UV-light and the band corresponding to the doubly labelled DNA strand was excised and pulverized. DNA was extracted by adding 1 mL of 1x TBE buffer and shaking at 4 °C for 2h. The gel debris was pelleted via centrifugation at 15000 rcf for 30 min (repeated once). The DNA was precipitated by addition of 1/10 volume of ammonium acetate solution (3 M, pH 5) and 2.5 volumes of ethanol. The sample was incubated at −80 °C for 1 h followed by a centrifugation step for 1 h at 4 °C. The supernatant was carefully decanted and the DNA was washed by addition of 5 mL of 70% ethanol and 30 min centrifugation at 15000 rcf. The supernatant was completely removed, the pellet dried for 10 min at 20 °C and resuspended in 10 mM Tris HCl pH 8.0 +50 mM NaCl.

### DNA origami preparation and purification

DNA origamis were assembled as described previously ^57^. In brief, scaffold DNA (25 nM), core staple strands (200 nM), force staple strands (400 nM), biotin adapter staple strands (200 nM) and the complementary doubly labelled promoter DNA strand (200 nM) were mixed in folding buffer (10 mM Tris pH 7.6, 1 mM EDTA, 20 mM MgCl_2_, 5 mM NaCl) and subjected to a multistep thermocycler protocol (**Supplementary Table 2**). Afterwards, the origami was purified by addition of one volume of 2x precipitation puffer (Tris HCl pH 7.6, 1 mM EDTA, 500 mM NaCl, 15% (w/v) PEG-8000) and centrifugation at 20000 rcf for 30 min at 4 °C. Afterwards, the supernatant was decanted and the pellet resuspended in 30 µL folding buffer for 30 min at 30 °C under constant shaking. All purification steps were repeated once.

### Restriction digestion of origami scaffolds

In order to generate force clamps with 0 pN force the spring strand was cleaved with a BamHI restriction endonuclease. To this end, 200 µM of the scaffold DNA and 3x molar excess of BamHI_comp strand were hybridized in FastDigest Green buffer (Thermo Scientific) by heating the sample to 90 °C followed by gradual cooling to 20 °C over 2 h. Afterwards, 1 U of FastDigest BamHI (Thermo Scientific) was added, incubated at 37 °C for 4 h. Subsequently, BamHI was heat inactivated at 80 °C for 10 min.

### Surface preparation

Silica microscope slides used for TIRF experiments were prepared as described before^52^. Briefly, fused silica slides (Plano) were cleaned in peroxomosulfuric acid (70% (v/v) sulfuric acid; Fisher Scientific, 10% (v/v) hydrogen peroxide; Sigma-Aldrich) for 30 min and washed with Millipore water under sonication. Afterwards, the slides were incubated in methanol for 20 min and sonicated for 5 min. For silan passivation, the slides were incubated in a freshly prepared N-[3-(Trimethoxisilyl)propyl]ethyldiamine (Sigma-Aldrich) solution (2% (v/v) in methanol with 4% (v/v) acetic acid) for 20 min, rinsed with methanol five times and an additional 20 times with Millipore water. The slides were dried for 1h at 37 °C. For polyethylene glycole (PEG) passivation, 100 µL of freshly prepared passivation solution (200 mg/mL methoxi-PEG succinimidyl valerate 5000 (Laysan Bio), 5 mg/mL biotin-PEG (Laysan Bio) in 1 mM NaHCO3) was sandwiched between a slide and a coverslip, incubated for 2 h and rinsed with Millipore water 20 times. The slides and coverslips were fully dried at 37 °C, vacuum-sealed in plastic tubes and stored at −20 °C.

### TIRF immobilisation assay

Single-molecule FRET measurements on immobilized DNA/protein complexes were carried out in custom-built flow-chambers based on fused silica slides passivated with polyethylene glycole (PEG). Flow chambers were prepared and assembled a described before ^44^.

For fluorescence measurements the flow chamber was incubated with 0.1 mg/mL NeutrAvidin (Pierce) in 1 x TBS (125 mM Tris/HCl pH 8, 150 mM NaCl) for 5 min and washed with 500 µL T78 buffer (100 mM Tris/HCl pH 7.8, 60 mM KCl, 5 mM MgCl_2_, 0.5 mg/mL BSA, 1% (v/v) glycerol). Afterwards, the chamber was flushed with DNA origami force clamps (10 pM in folding buffer) for 5 s and washed with 500 µL T78 buffer. The chamber wash flushed with photostabilizer buffer (T78 buffer with 2 mM Trolox, 1% (w/v) D-glucose, 7.5 U/mL glucose oxidase type VII (Sigma Aldrich), 1 kU/mL catalase (Sigma Aldrich)) supplemented with 10 nM human TBP and incubated for 5 min before starting video acquisition.

### Wide-field single-molecule detection and data analysis

Time resolved single-molecule fluorescence measurements were performed on a homebuilt prism-type total internal reflection setup based on a Leica DMi8 inverse research microscope. Fluorophores were exited with a 532 nm solid state laser (Coherent OBIS) with a power of 30 mW and 637 nm diode laser (Coherent OBIS, clean-up filter ZET 635/10, AHF Göttingen) with a power of 50 mW employing alternating laser excitation (Multistream, Cairn Reasearch, UK) ^75^. The fluorescence was collected by a Leica HC PL Apo 63x N.A. 1.20 water-immersion objective and split by wavelength with a dichroic mirror (HC BS 640, AHF) into two detection channels that were further filtered with a 582/75 bandpass filter (Brightline HC, AHF) in the green channel and a 635 nm long-pass filter (LP Edge Basic, AHF) in the red detection channel. Both detection channels were recorded by one EMCCD camera (Andor IXon Ultra 897, EM-gain 20, framerate 40 Hz, 400 frames) in a dual-view configuration (TripleSplit, Cairn Research).

The videos were analysed employing the iSMS software ^76^ using the programs defaults settings. Molecule spots were detected using a threshold of 100 for ATTO 532 and ATTO 647N spots. FRET efficiencies were calculated as proximity ratios from fluorescence intensity time traces that were corrected for background fluorescence using the average intensity of all pixels with a 2 pixel distance to the molecule spot. For TBP dwell time histograms, traces showing dynamic switching between FRET states were fitted with the vbFRET algorithm ^77^ limited to two states.

FRET efficiency histograms were calculated from all frames of traces showing dynamic switching between states with an S-value between 0.4 to 0.6 and were fitted with a Gaussian distribution. All states calculated with vbFRET with a FRET efficiency within the FWHM of a fitted FRET population were used to calculate the dwell time histogram. The histograms of at least three independent experiments were normalized and fitted with a monoexponential decay function to calculate the mean dwell time in the high FRET state (TBP bound to DNA).

### Confocal Single-pair FRET measurements

Prior to sample loading, the sample chambers (Cellview slide, Greiner Bio-One) were passivated with 10 mM Tris/HCl pH 8 with 2 mg/mL BSA for 10 min and washed once with T78 buffer.

For equilibrium measurements (**Figure 2, Figure 3, Supplementary Figure 4**) complexes were formed with 20 pM DNA origami and 20 nM TBP, Brf2 and Bdp1 or 200 nM TFIIB and incubated for 30 min at room temperature in T78 buffer with 2 mM DTT. For time course experiments (**Figure 4, Supplementary Figure 6**) 20 pM DNA origami and 20 nM Brf2 and Bdp1 or 200 nM of TFIIB in T78 buffer with 2 mM DTT were added to the sample chamber and data acquisition was started to measure the unbound DNA state. After 2 minutes, TBP was added to initiate complex formation.

Single-molecule fluorescence of diffusing complexes was detected with a MicroTime 200 confocal microscope (Picoquant) equipped with pulsed laser diodes (532 nm: LDH-P-FA-530B; 636 nm: LDH-D-C-640; PicoQuant / cleanup filter: zet635; Chroma). The fluorophors were excited at 20 µW using pulsed interleaved excitation. Emitted fluorescence was collected using a 1.2 NA, ×60 microscope objective (UplanSApo ×60/1.20W; Olympus) and a 50-μm confocal pinhole. A dichroic mirror (T635lpxr; Chroma) separated donor and acceptor fluorescence. Additional bandpass filters (donor: ff01-582/64; Chroma; acceptor: H690/70; Chroma) completed spectral separation of the sample fluorescence. Each filtered photon stream was detected by an individual APD (SPCM-AQRH-14-TR, Exceliatas Technologies) and analyzed by a TCSPC capable PicoQuant HydraHarp 400.

### Data analysis

Data analysis of confocal FRET measurements was performed with the software package PAM^78^. Photon bursts of diffusing molecules were determined by an all-photon burst search (APBS, parameters: *L*=50, M=20, and *T*=500 μs) and an additional dual-channel burst search (DCBS, parameters: *L*=50, *M*_*GG+GR*_=20, M_RR_=20, and *T*=500 μs). Burst data were corrected for donor leakage and direct excitation of the acceptor (determined from APBS according to ^79^) as well as γ and β (determined from DCBS ES-histograms using an internal fit on multiple E/S separated FRET populations). The data were binned (bin size =0.025), plotted as E histogram and fitted with a single (DNA) or triple Gaussian fit.

### Kinetics measurements

Data were processed as above. All bursts were sorted according to their FRET efficiency (low FRET for E<0.3 and high FRET for E>0.6) and binned by macrotime (bin size=2 min). Low FRET and high FRET bins were normalized to the combined sum to determine relative ratios of both populations which were plotted against time and fitted with a mono-exponential function. The fit-derived decay constant and y-offset (y0, equivalent to low FRET ratio at equilibrium) for the low FRET population were used to determine dwell times in the high FRET and low FRET state via deconvolution with a perturbation-relaxation model (see also **Supplementary informations**).

## Acknowledgements

We gratefully acknowledge financial support by the Deutsche Forschungsgemeinschaft (SFB960-TP7 to D.G.). PT acknowledges support by the DFG (grant INST 86/1904-1 FUGG), excellence clusters CIPSM (Center for Integrated Protein Science Munich) and NIM (Nanosystems Initiative Munich). TL is also supported through NIM and the SFB1032-TP6. FH and CE were supported by DFGs ‘Emmy-Noether-Programm’ [DFG grant no. EN 1204/1-1] and and SFB960-TP A8. A.V. is supported by a Cancer Research UK Programme Foundation (CR-UK C47547/A21536) and a Wellcome Trust Investigator Award (200818/Z/16/Z).

Furthermore, we would like to thank Dr. Sarah Willkomm for advice on analysing the kinetics data, Michael Pilsl for support on the EM measurements and Elisabeth Piechatschek and Elke Papst for technical assistance.

## Author contributions

D.G. and K.K. conceived the study. K.K. performed the single-molecule measurements. K.K. and T.S. analyzed the single-molecule data. J.G. and A.V. purified the proteins. T.S., A.M.V., T. L. and P. T. designed and manufactured the DNA origami force clamp. F.H. and C.E. carried out electron microscopy measurements and analysed the data. K.K. and D.G. wrote the paper. All authors commented on the paper.

## References

1. Werner, F. & Grohmann, D. Evolution of multisubunit RNA polymerases in the three domains of life. Nature reviews. Microbiology 9, 85–98; 10.1038/nrmicro2507 (2011).

2. Dienemann, C., Schwalb, B., Schilbach, S. & Cramer, P. Promoter Distortion and Opening in the RNA Polymerase II Cleft. Molecular cell 73, 97–106.e4; 10.1016/j.molcel.2018.10.014 (2019).

3. Griesenbeck, J., Tschochner, H. & Grohmann, D. Structure and Function of RNA Polymerases and the Transcription Machineries. Sub-cellular biochemistry 83, 225–270; 10.1007/978-3-319-46503-6_9 (2017).

4. Liu, X., Bushnell, D. A., Wang, D., Calero, G. & Kornberg, R. D. Structure of an RNA polymerase II-TFIIB complex and the transcription initiation mechanism. Science (New York, N.Y.) 327, 206–209; 10.1126/science.1182015 (2010).

5. Sainsbury, S., Niesser, J. & Cramer, P. Structure and function of the initially transcribing RNA polymerase II–TFIIB complex. Nature 493, 437–440; 10.1038/nature11715 (2012).

6. Kramm, K., Engel, C. & Grohmann, D. Transcription initiation factor TBP: old friend new questions. Biochemical Society Transactions 47, 411–423; 10.1042/BST20180623 (2019).

7. Vannini, A. & Cramer, P. Conservation between the RNA polymerase I, II, and III transcription initiation machineries. Molecular cell 45, 439–446; 10.1016/j.molcel.2012.01.023 (2012).

8. Engel, C., Neyer, S. & Cramer, P. Distinct Mechanisms of Transcription Initiation by RNA Polymerases I and II. Annual review of biophysics 47, 425–446; 10.1146/annurev-biophys-070317-033058 (2018).

9. Basehoar, A. D., Zanton, S. J. & Pugh, B.F. Identification and Distinct Regulation of Yeast TATA Box-Containing Genes. Cell 116, 699–709; 10.1016/S0092-8674(04)00205-3 (2004).

10. Savinkova, L. et al. An Experimental Verification of the Predicted Effects of Promoter TATA-Box Polymorphisms Associated with Human Diseases on Interactions between the TATA Boxes and TATA-Binding Protein. PLoS ONE 8, e54626; 10.1371/journal.pone.0054626 (2013).

11. Wobbe, C. R. & Struhl, K. Yeast and human TATA-binding proteins have nearly identical DNA sequence requirements for transcription in vitro. Molecular and cellular biology 10, 3859–3867; 10.1128/MCB.10.8.3859.Updated (1990).

12. Smollett, K., Blombach, F., Reichelt, R., Thomm, M. & Werner, F. A global analysis of transcription reveals two modes of Spt4/5 recruitment to archaeal RNA polymerase. Nature Microbiology 2, 17021; 10.1038/nmicrobiol.2017.21 (2017).

13. Krebs, A. R. et al. Genome-wide Single-Molecule Footprinting Reveals High RNA Polymerase II Turnover at Paused Promoters. Molecular cell 67, 411 – 422.e4; 10.1016/J.MOLCEL.2017.06.027 (2017).

14. Kim, T. H. et al. A high-resolution map of active promoters in the human genome. Nature 436, 876–880; 10.1038/nature03877 (2005).

15. Kim, J. L., Nikolov, D. B. & Burley, S. K. Co-crystal structure of TBP recognizing the minor groove of a TATA element. Nature 365, 520–527; 10.1038/365520a0 (1993).

16. Kim, Y., Geiger, J. H., Hahn, S. & Sigler, P. B. Crystal structure of a yeast TBP/TATA-box complex. Nature 365, 512–520; 10.1038/365512a0 (1993).

17. Kosa, P. F., Ghosh, G., DeDecker, B. S. & Sigler, P. B. The 2.1-A crystal structure of an archaeal preinitiation complex: TATA-box-binding protein/transcription factor (II)B core/TATA-box. Proceedings of the National Academy of Sciences of the United States of America 94, 6042–6047; 10.1073/PNAS.94.12.6042 (1997).

18. Kostrewa, D. et al. RNA polymerase II–TFIIB structure and mechanism of transcription initiation. Nature 462, 323–330; 10.1038/nature08548 (2009).

19. Hantsche, M. & Cramer, P. Conserved RNA polymerase II initiation complex structure. Current Opinion in Structural Biology 47, 17–22; 10.1016/j.sbi.2017.03.013 (2017).

20. He, Y. et al. Near-atomic resolution visualization of human transcription promoter opening. Nature 533, 359–365; 10.1038/nature17970 (2016).

21. Abascal-Palacios, G., Ramsay, E. P., Beuron, F., Morris, E. & Vannini, A. Structural basis of RNA polymerase III transcription initiation. Nature 553, 301–306; 10.1038/nature25441 (2018).

22. Sadian, Y. et al. Structural insights into transcription initiation by yeast RNA polymerase I. The EMBO Journal 36, 2698–2709; 10.15252/embj.201796958 (2017).

23. Vorländer, M. K., Khatter, H., Wetzel, R., Hagen, W. J.H. H. & Müller, C. W. Molecular mechanism of promoter opening by RNA polymerase III. Nature 553, 295–300; 10.1038/nature25440 (2018).

24. Sainsbury, S., Bernecky, C. & Cramer, P. Structural basis of transcription initiation by RNA polymerase II. Nature Reviews Molecular Cell Biology 16, 129–143; 10.1038/nrm3952 (2015).

25. Hausner, W., Wettach, J., Hethke, C. & Thomm, M. Two transcription factors related with the eucaryal transcription factors TATA-binding protein and transcription factor IIB direct promoter recognition by an archaeal RNA polymerase. The Journal of biological chemistry 271, 30144–30148; 10.1074/JBC.271.47.30144 (1996).

26. Werner, F. & Weinzierl, R. O. J. A recombinant RNA polymerase II-like enzyme capable of promoter-specific transcription. Molecular cell 10, 635–646 (2002).

27. Buratowski, S., Hahn, S., Guarente, L. & Sharp, P. A. Five intermediate complexes in transcription initiation by RNA polymerase II. Cell 56, 549–561; 10.1016/0092-8674(89)90578-3 (1989).

28. Cavallini, B., Huet, J., Plassat, J. L., Sentenac, A. & Egly, J. M. A yeast activity can substitute for the HeLa cell TATA box factor. Nature 334, 77–80; 10.1038/334077a0 (1988).

29. Bell, S. D. & Jackson, S. P. The Role of Transcription Factor B in Transcription Initiation and Promoter Clearance in the Archaeon Sulfolobus acidocaldarius. Journal of Biological Chemistry 275, 12934–12940; 10.1074/jbc.275.17.12934 (2000).

30. Khatter, H., Vorländer, M. K. & Müller, C. W. RNA polymerase I and III: similar yet unique. Current Opinion in Structural Biology 47, 88–94; 10.1016/j.sbi.2017.05.008 (2017).

31. Eberhard, D., Tora, L., marc Egly, J. & Grummt, I. A TBP-containing multiprotein complex (TIF-IB) mediates transcription specificity of murine RNA polymerase I. Nucleic Acids Research 21, 4180–4186; 10.1093/nar/21.18.4180 (1993).

32. Steffan, J. S., Keys, D. A., Dodd, J. A. & Nomura, M. The role of TBP in rDNA transcription by RNA polymerase I in Saccharomyces cerevisiae: TBP is required for upstream activation factor-dependent recruitment of core factor. Genes and Development 10, 2551–2563; 10.1101/gad.10.20.2551 (1996).

33. Siddiqi, I., Keener, J., Vu, L. & Nomura, M. Role of TATA binding protein (TBP) in yeast ribosomal dna transcription by RNA polymerase I: defects in the dual functions of transcription factor UAF cannot be suppressed by TBP. Molecular and cellular biology 21, 2292–2297; 10.1128/MCB.21.7.2292-2297.2001 (2001).

34. Keener, J., Dodd, J. A., Lalo, D. & Nomura, M. Histones H3 and H4 are components of upstream activation factor required for the high-level transcription of yeast rDNA by RNA polymerase I. Proceedings of the National Academy of Sciences of the United States of America 94, 13458–13462; 10.1073/PNAS.94.25.13458 (1997).

35. Bedwell, G. J., Appling, F. D., Anderson, S. J. & Schneider, D. A. Efficient transcription by RNA polymerase I using recombinant core factor. Gene 492, 94–99; 10.1016/j.gene.2011.10.049 (2012).

36. Keener, J., Josaitis, C. A., Dodd, J. A. & Nomura, M. Reconstitution of yeast RNA polymerase I transcription in vitro from purified components: TATA-binding protein is not required for basal transcription. Journal of Biological Chemistry 273, 33795–33802; 10.1074/jbc.273.50.33795 (1998).

37. Schramm, L. & Hernandez, N. Recruitment of RNA polymerase III to its target promoters. Genes & Development 16, 2593–2620; 10.1101/gad.1018902 (2002).

38. Kassavetis, G. A. A minimal RNA polymerase III transcription system. The EMBO Journal 18, 5042–5051; 10.1093/emboj/18.18.5042 (1999).

39. Verma, N. et al. Bdp1 interacts with SNAPc bound to a U6, but not U1, snRNA gene promoter element to establish a stable protein-DNA complex. FEBS Letters 592, 2489–2498; 10.1002/1873-3468.13169 (2018).

40. Kassavetis, G. A. & Geiduschek, E. P. Transcription factor TFIIIB and transcription by RNA polymerase III. Biochemical Society Transactions 34, 1082–1087; 10.1042/BST0341082 (2006).

41. Schramm, L., Pendergrast, P. S., Sun, Y. & Hernandez, N. Different human TFIIIB activities direct RNA polymerase III transcription from TATA-containing and TATA-less promoters. Genes and Development 14, 2650–2663; 10.1101/gad.836400 (2000).

42. Kassavetis, G. A., Braun, B. R., Nguyen, L. H. & Geiduschek, E. P. S. cerevisiae TFIIIB is the transcription initiation factor proper of RNA polymerase III, while TFIIIA and TFIIIC are assembly factors. Cell 60, 235–245 (1990).

43. Blair, R. H., Goodrich, J. A. & Kugel, J. F. Single-molecule fluorescence resonance energy transfer shows uniformity in TATA binding protein-induced DNA bending and heterogeneity in bending kinetics. Biochemistry 51, 7444–7455; 10.1021/bi300491j (2012).

44. Gietl, A. et al. Eukaryotic and archaeal TBP and TFB/TF(II)B follow different promoter DNA bending pathways. Nucleic Acids Research 42, 6219–6231; 10.1093/nar/gku273 (2014).

45. Schluesche, P. et al. Dynamics of TBP binding to the TATA box. 16057422 6862, 68620E–68620E-8; 10.1117/12.769177 (2008).

46. Heiss, G. et al. Conformational changes and catalytic inefficiency associated with Mot1-mediated TBP-DNA dissociation. Nucleic Acids Research 47, 2793–2806; 10.1093/nar/gky1322 (2019).

47. Zarrabi, N., Schluesche, P., Meisterernst, M., Börsch, M. & Lamb, D. C. Analyzing the Dynamics of Single TBP-DNA-NC2 Complexes Using Hidden Markov Models. Biophysical journal 115, 2310–2326; 10.1016/j.bpj.2018.11.015 (2018).

48. Delgadillo, R. F., Whittington, J. D. E., Parkhurst, L. K. & Parkhurst, L. J. The TATA-binding protein core domain in solution variably bends TATA sequences via a three-step binding mechanism. Biochemistry 48, 1801–1809; 10.1021/bi8018724 (2009).

49. Masters, K. M., Parkhurst, K. M., Daugherty, M. A. & Parkhurst, L. J. Native human TATA-binding protein simultaneously binds and bends promoter DNA without a slow isomerization step or TFIIB requirement. Journal of Biological Chemistry 278, 31685–31690; 10.1074/jbc.M305201200 (2003).

50. Parkhurst, K. M., Richards, R. M., Brenowitz, M. & Parkhurst, L. J. Intermediate species possessing bent DNA are present along the pathway to formation of a final TBP-TATA complex 1 1Edited by R. Ebright. Journal of Molecular Biology 289, 1327–1341; 10.1006/jmbi.1999.2835 (1999).

51. Whittington, J. D. E. et al. TATA-binding protein recognition and bending of a consensus promoter are protein species dependent. Biochemistry 47, 7264–7273; 10.1021/bi800139w (2008).

52. Gouge, J. et al. Molecular mechanisms of Bdp1 in TFIIIB assembly and RNA polymerase III transcription initiation. Nature Communications 8, 130; 10.1038/s41467-017-00126-1 (2017).

53. Tyagi, M., Imam, N., Verma, K. & Patel, A. K. Chromatin remodelers: We are the drivers!! Nucleus (Austin, Tex.) 7, 388–404; 10.1080/19491034.2016.1211217 (2016).

54. Struhl, K. & Segal, E. Determinants of nucleosome positioning. Nature Structural & Molecular Biology 20, 267–273; 10.1038/nsmb.2506 (2013).

55. Funke, J. J. et al. Uncovering the forces between nucleosomes using DNA origami. Science advances 2, e1600974; 10.1126/sciadv.1600974 (2016).

56. Yan, J. & Marko, J. F. Effects of DNA-distorting proteins on DNA elastic response. Physical review. E, Statistical, nonlinear, and soft matter physics 68, 11905; 10.1103/PhysRevE.68.011905 (2003).

57. Nickels, P. C. et al. Molecular force spectroscopy with a DNA origami-based nanoscopic force clamp. Science (New York, N.Y.) 354, 305–307; 10.1126/science.aah5974 (2016).

58. Smith, S. B., Cui, Y. & Bustamante, C. Overstretching B-DNA: the elastic response of individual double-stranded and single-stranded DNA molecules. Science 271, 795–799 (1996).

59. Kouzine, F., Levens, D. & Baranello, L. DNA topology and transcription. Nucleus (Austin, Tex.) 5, 195–202; 10.4161/nucl.28909 (2014).

60. Gietl, A. & Grohmann, D. Modern biophysical approaches probe transcription-factor-induced DNA bending and looping. Biochemical Society Transactions 41, 368–373; 10.1042/BST20120301 (2013).

61. Chien, F.-T. & van der Heijden, T. Characterization of nucleosome unwrapping within chromatin fibers using magnetic tweezers. Biophysical journal 107, 373–383; 10.1016/j.bpj.2014.05.036 (2014).

62. Kruithof, M. et al. Single-molecule force spectroscopy reveals a highly compliant helical folding for the 30-nm chromatin fiber. Nature Structural & Molecular Biology 16, 534–540; 10.1038/nsmb.1590 (2009).

63. Cui, Y. & Bustamante, C. Pulling a single chromatin fiber reveals the forces that maintain its higher-order structure. Proceedings of the National Academy of Sciences of the United States of America 97, 127–132; 10.1073/pnas.97.1.127 (2000).

64. Meng, H., Andresen, K. & van Noort, J. Quantitative analysis of single-molecule force spectroscopy on folded chromatin fibers. Nucleic Acids Research 43, 3578–3590; 10.1093/nar/gkv215 (2015).

65. Dieci, G. & Sentenac, A. Facilitated recycling pathway for RNA polymerase III. Cell 84, 245–252 (1996).

66. Dieci, G., Bosio, M. C., Fermi, B. & Ferrari, R. Transcription reinitiation by RNA polymerase III. Biochimica et biophysica acta 1829, 331–341; 10.1016/j.bbagrm.2012.10.009 (2013).

67. Dieci, G. & Sentenac, A. Detours and shortcuts to transcription reinitiation. Trends in Biochemical Sciences 28, 202–209; 10.1016/S0968-0004(03)00054-9 (2003).

68. Han, Y., Yan, C., Fishbain, S., Ivanov, I. & He, Y. Structural visualization of RNA polymerase III transcription machineries. Cell Discovery 4, 40; 10.1038/s41421-018-0044-z (2018).

69. Helbo, A. S., Lay, F. D., Jones, P. A., Liang, G. & Grønbæk, K. Nucleosome Positioning and NDR Structure at RNA Polymerase III Promoters. Scientific Reports 7, 41947; 10.1038/srep41947.

70. Schones, D. E. et al. Dynamic regulation of nucleosome positioning in the human genome. Cell 132, 887–898; 10.1016/j.cell.2008.02.022 (2008).

71. Zhang, Y. et al. Intrinsic histone-DNA interactions are not the major determinant of nucleosome positions in vivo. Nature Structural & Molecular Biology 16, 847–852; 10.1038/nsmb.1636 (2009).

72. Radman-Livaja, M. & Rando, O. J. Nucleosome positioning: how is it established, and why does it matter? Developmental biology 339, 258–266; 10.1016/j.ydbio.2009.06.012 (2010).

73. Sambrook, J. Molecular cloning. 3rd ed. (Cold Spring Harbor Laboratory Press, Cold Spring Harbor, NY, 2001).

74. Douglas, S. M., Chou, J. J. & Shih, W. M. DNA-nanotube-induced alignment of membrane proteins for NMR structure determination. Proceedings of the National Academy of Sciences of the United States of America 104, 6644–6648; 10.1073/pnas.0700930104 (2007).

75. Kapanidis, A. N. et al. Alternating-laser excitation of single molecules. Accounts of chemical research 38, 523–533; 10.1021/ar0401348 (2005).

76. Preus, S., Noer, S. L., Hildebrandt, L. L., Gudnason, D. & Birkedal, V. iSMS: single-molecule FRET microscopy software. Nature Methods 12, 593; 10.1038/nmeth.3435 (2015).

77. Bronson, J. E., Fei, J., Hofman, J. M., Gonzalez, R. L. & Wiggins, C. H. Learning rates and states from biophysical time series: a Bayesian approach to model selection and single-molecule FRET data. Biophysical journal 97, 3196–3205; 10.1016/j.bpj.2009.09.031 (2009).

78. Schrimpf, W., Barth, A., Hendrix, J. & Lamb, D. C. PAM: A Framework for Integrated Analysis of Imaging, Single-Molecule, and Ensemble Fluorescence Data. Biophysical journal 114, 1518–1528; 10.1016/j.bpj.2018.02.035 (2018).

79. Hellenkamp, B. et al. Precision and accuracy of single-molecule FRET measurements-a multi-laboratory benchmark study. Nature Methods 15, 669–676; 10.1038/s41592-018-0085-0 (2018).

