## Supplemental data for "DNA origami-based single-molecule force spectroscopy unravels the molecular basis of RNA Polymerase III pre-initiation complex stability"

### Supplementary material and methods

**Supplementary Table 1: Sequences of DNA oligonucleotides**. The coupling position of the Atto532 (green) and Atto647N (red) fluorophore, the TATA box sequence (blue) and the BamHI restriction site (yellow) are indicated by color. The 5'-phosphate (Pho-) modification was used for enzymatic ligation (see below).

| Name | Sequences and modifications of DNA oligonucleotides (5'-3') | Supplier |
|---|---|---|
| AdMLP forward | Pho-gatccTATGACTGCTTCGCGCCCGAACCCCCTTTTATAGGCGCCCTAa | Biomers |
| AdMLP reverse | Pho-agcttTAGGGCGCCTATAAAAGGGGGTTCGGGCGCGAAGCAGTCATAg | Biomers |
| U6 forward | Pho-gatccTATGATCAAGGGTTACTCTAACACCTATTTTAAGCCCTTCAATCAAATCATCTTGGTCCGa | Biomers |
| U6 reverse | Pho-agcttCGGACCAAGATGATTTGATTGAAGGGCTTAAAATAGGTGTTAGAGTAACCCTTGATCATAg | Biomers |
| MLP_D | AAGCTTTAGGGCGCCTATAAAAG | IBA |
| MLP_A | Pho-GGGGTTCGGGCG | IBA |
| MLP_lig | TTCGCGCCCGAACCCCCTTTTATAGGCGC | Sigma |
| MLP_TS | CGCCCGAACCCCCTTTTATAGGCGCCCTAAAGCTT | MWG |
| U6_D | CGGACCAAGATGATTTGATTGAAGGGCTTAAAATA | IBA |
| U6_A | Pho-GGTGTTAGAGTAACCCTTGA | IBA |
| U6_Lig | GGTTACTCTAACACCTATTTTAAGCCCTTC | Sigma |
| U6_TS | TCAAGGGTTACTCTCACACCTATTTTAAGCCCTTCAATCAAATCATCTTGGTCCG | Sigma |
| BamHI_comp | TCATAGGATCCCCGGGTA | Metabion |
| MLP_comp | AAGCTTTAGGGCGCCTATAAAAGGGGGTTCGGGCG | see below |
| U6_comp | CGGACCAAGATGATTTGATTGAAGGGCTTAAAATAGGTGTTAGAGTAACCCTTGA | see below |
| AdMLP 6pN_1 | TTTTGCTTTCATCAACATTAAATCCGTAATCGTAACCTTGGGTACAGG | MWG |
| AdMLP 6pN_2 | TTTTAAGTTTCATTCCTCTGGAGAGGCTATACGCCAGGGTTTTCCCAGT | MWG |
| AdMLP 6pN_3 | TTTTCGCTCATGGACGAGCCG | MWG |
| AdMLP 6pN_4 | TTTTACTTGCCTGAGTAGAAGTAATTCCACGATT | MWG |
| AdMLP 6pN_5 | AGCCTTCACCGCCTGGCGTTATCCGCTCACATAAC | MWG |
| AdMLP 6pN_6 | GCTACAACATAAATACCATTGCAACAGGAAAAATTTT | MWG |
| AdMLP 6pN_7 | ATGCAATGGTGAGAAAGGCATGATTAAGGTGCATCAGATTGTAATT | MWG |
| AdMLP 6pN_8 | GTCAATAGCAAGGCACAGGCACCTCAGAGCTTTAA | MWG |
| AdMLP 6pN_9 | TTTCCTGTGTGAAATTCCTGAGAGGGGTCG | MWG |
| AdMLP 6pN_10 | GGGGATGTGCTACCTGTTTAGCTTTTT | MWG |

**Supplementary Table 2: Temperature ramp for the folding of DNA origami force clamps**. The DNA origami mix including all staples was heated for 2 min (lid temperature 95°C) and subsequently annealed using a non-linear temperature ramp.

| Temperature (°C) | Time per °C (min) | Temperature (°C) | Time per °C (min) |
|---:|---:|---:|---:|
| 65 | 2 | 44 | 75 |
| 64-61 | 3 | 43 | 60 |
| 60-59 | 15 | 42 | 45 |
| 58 | 30 | 41-39 | 30 |
| 57 | 45 | 38-37 | 15 |
| 56 | 60 | 36-30 | 8 |
| 55 | 75 | 29-25 | 2 |
| 54-45 | 90 | 8 | storage |

### Calculation of forces for the AdML promoter and U6 promoter DNA origami force clamps

Forces were calculated according to reference [1] where the ssDNA is described as a freely-jointed chain (FJC). Width of the double-stranded promotor is the average value of the bent and unbent state. For the AdML promotor we calculated an average width of 10.17 nm and for the U6 promotor of 16.01 nm. For new scaffold and promotor length the forces for the 2.5 pN staple set form reference [1] are calculated to be 3.3 pN for the AdML promotor and 2.6 pN for the U6 promotor. The 6.2 pN staple set of reference [1] results in a 6.6 pN force for the U6 promotor. The 6.0 pN for the AdML promotor were new designed. Staples were ordered from Eurofins MWG and are listed in **Supplementary Table 1**.

### Sample preparation and transmission electron microscopy imaging of DNA origami force clamps

Electron microscopy grids (copper, 400 mesh; PlanoEM, Germany) were pre-coated with a ~8 nm thick carbon support film generated with a 'Turbo Carbon Coater' (Cressington Scientific Instruments, UK). Four microliters of force clamp solution (10 nM) was applied to glow-discharged grids for 30 s, blotted off and the grids washed for 30 s (bi-distilled water). Grids were dried and imaged without staining using a JEM2100F transmission electron microscope (Jeol, Japan) equipped with TemCamF416 detector (TVIPS, Germany). A total of 94 micrographs were semi-automatically collected at 40,000 magnification (2.7 Å/pix) and a defocus range of -2.5 to -4.5 µm using the Serial-EM software package [2]. CTF-estimation and manual particle picking were carried out in Relion 3 [3]. A total of 2,133 particles (2x binned; box size 180 pixels/100nm) were extracted, contrast-inverted and 2-D-classified into three classes with CTF-amplitude correction from the first peak onward.

#### Fluorescence correlation spectroscopy

FCS analysis was performed with the PicoQuant software package SymPhoTime 64. The acceptor signal of an arbitrary 10 min interval of each data set was autocorrelated according to eq. 1 and fitted following eq. 2 with one (dsDNA) or two (origami) diffusion components ($n_{\mathrm{Diff}}$). The relative diffusion times given in **Supplementary Figure 3** were calculated with a non-calibrated confocal volume $V_{\mathrm{eff}}=1$. The statistical error shown is the standard error of the fit.

$$G(\tau) = \frac{\langle I(t)\, I(t+\tau)\rangle}{\langle I(t)\rangle^2} - 1 \qquad \text{(eq. 1)}$$

with:

$\tau$: correlation time

$I$: signal intensity

$t$: experiment time

$$G(t) = \left[1 + T\left[\exp\left(\frac{t}{\tau_{Trip}}\right) - 1\right]\right] \sum_{i=0}^{n_{Diff}-1} \frac{\rho[i]}{\left[1+\frac{t}{\tau_{Diff}[i]}\right]\left[1+\frac{t}{\tau_{Diff}[i]\kappa^2}\right]^{0.5}} \qquad \text{(eq. 2)}$$

with:

$t$: correlation time

$T$: fraction of triplet state molecules

$\tau_{\mathrm{Trip}}$: lifetime of the triplet state

$\rho$: contribution of the $i^{\mathrm{th}}$ diffusing species

$\kappa$: length to diameter ratio of the confocal volume

#### Confocal kinetics measurements

The unbent and bent state dwell times for the highly stable AdMLP-TBP-(TFIIB) and U6-TBP-Brf2-(Bdp1) complexes were measured via confocal single-molecule experiments in solution. All factors except TBP were mixed prior to data acquisition and TBP was added 2 min after the start and the decay of the low FRET and increase of the high FRET population was monitored over time. Fitting this data to a mono-exponential model yields the decay rate $\tau$ and the unbent fraction $[u]_{new}$ in equilibrium with all proteins. This decay constant translates to the complex assembly rate that is superimposed with the simultaneous complex disassembly. We used a perturbation-relaxation kinetics model to extract both rates from the data.

The DNA-TBP-system is in an equilibrium of two states. The unbent state and the bent state.

$$\text{unbent} \underset{k_{unbent} = \tau_{bent}^{-1}}{\overset{k_{bent} = \tau_{unbent}^{-1}}{\rightleftharpoons}} \text{bent}$$

Where $k_{bent}$ is the bending rate and $k_{unbent}$ is the unbending rate. Conversely, $\tau_{unbent}$ and $\tau_{bent}$ are the average lifetimes of a molecule in the unbent or bent state. The equilibrium constant K is given by the ratio of the rates:

$$K = \frac{k_{bent}}{k_{unbent}}$$

The system is perturbed by a change in the TBP concentration and will relax in a first order process to its new equilibrium since $k_{bent}$ is concentration dependent. The unbent fraction $[u]$ will vary according to

$$[u](t) = [u]_{new} + Ae^{-\frac{t}{\tau}}$$

The amplitude $A$ is the change of the unbent fraction and. The new equilibrium unbent fraction is $[u]_{new}$. The relaxation time $\tau$ is given by both rate constants.

$$\tau = \frac{1}{k_{bent} + k_{unbent}}$$

With the new equilibrium constant and the decay time both rate constants can be calculated for the new equilibrium.

$$k_{bent} = \frac{K}{\tau \cdot (K+1)} = \frac{1}{\tau_{unbent}}$$

$$k_{unbent} = \frac{1}{\tau \cdot (K+1)} = \frac{1}{\tau_{bent}}$$

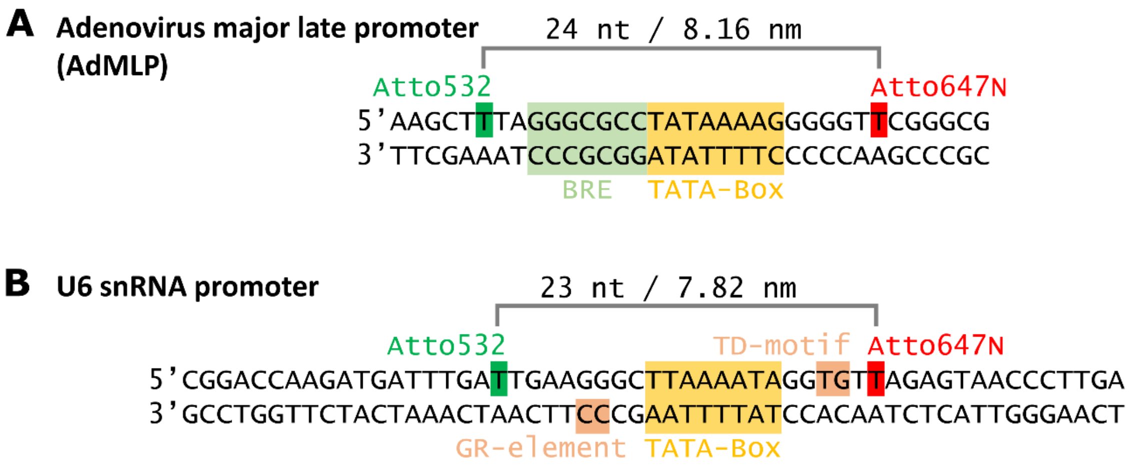


**Supplementary figure 1: Schematic overview of the used promoter DNA sequences. A)** DNA sequence of the Adenovirus major late promoter (AdMLP). The TATA-box element (yellow), bound by TBP, the B recognition element (BRE, light green) bound by TFIIB and the position of the donor (green) and acceptor fluorophore (red) are indicated by colour. **B)** DNA sequence of the U6 snRNA promoter. The TATA-box element (yellow), bound by TBP, the GR-element and TD-motif (orange) bound by Brf2 and the position of the donor (green) and acceptor fluorophore (red) are indicated by colour.

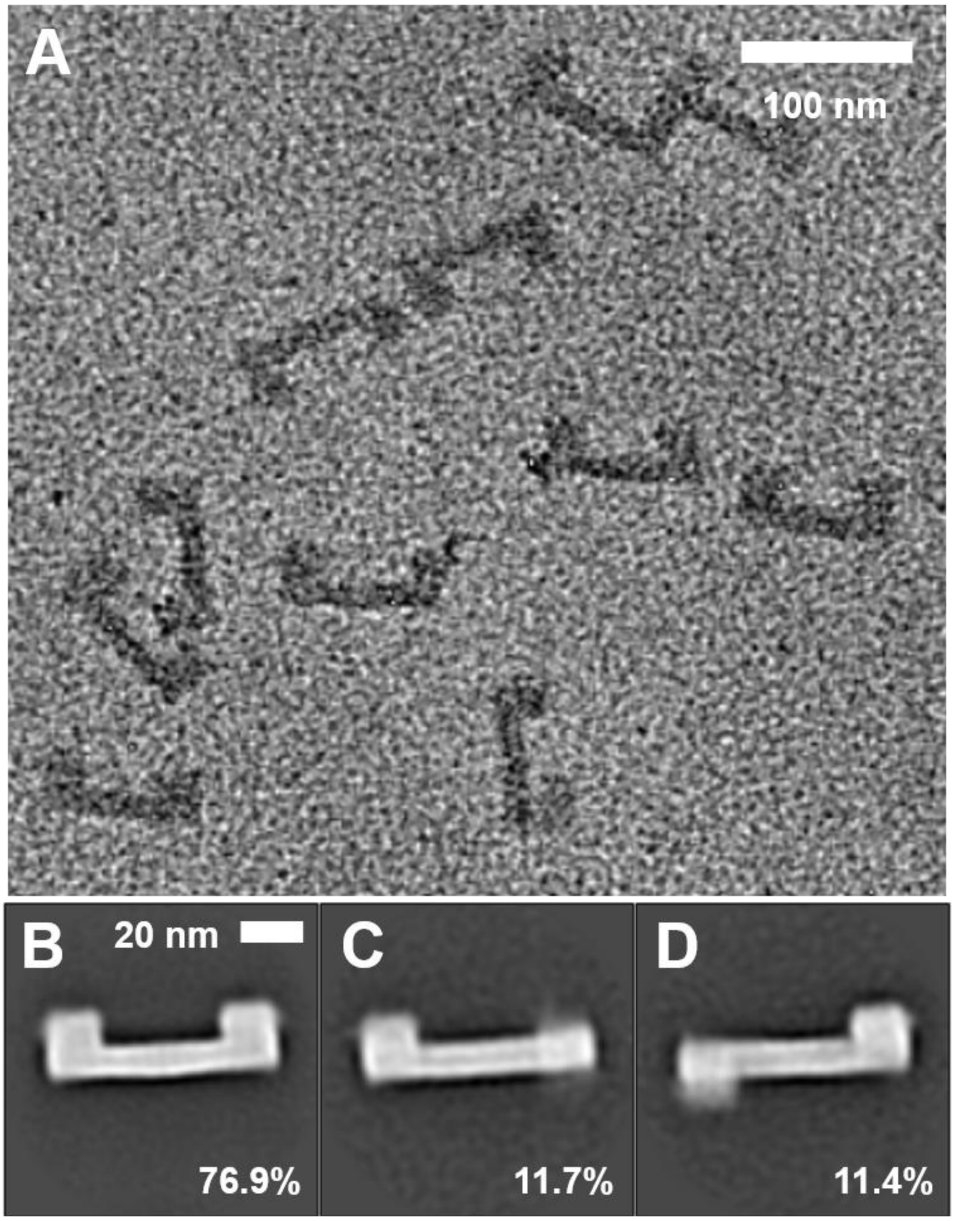


**Supplementary Figure 2: Electron Microscopy characterization 3pN DNA origami force clamps. A**) Exemplary section of electron micrograph showing unstained 'force clamps'. Scale bar: 100 nm. **B**) 2D-class average 1 (Methods) comprises 76.9% of particles and shows mainly intact origamis with limited flexibility in peripheral regions as previously observed [1]. Scale bar: 20 nm. **C**) 2D-class average 2 comprises 11.7% of particles. Unstained particles in this class appear to be damaged either during origami assembly or grid-preparation. Scale bar: 20 nm. **D**) 2D-class average 3 comprises 11.4% of particles. Particles in this class may be in top-view and/or could be damaged.

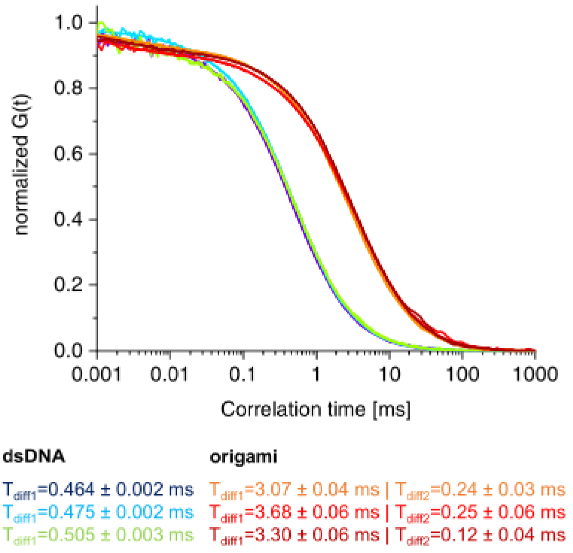


**Supplementary Figure 3. Fluorescence correlation spectroscopy monitors diffusion behaviour of a DNA origami force clamp compared to the respective short double-stranded promoter DNA**. The autocorrelation function G(t) of the acceptor signal of the short dsDNA U6 promoter (55 nt) and the U6 DNA origami force clamp were calculated. The decay was fitted with a one- (dsDNA) or two-component (DNA origami force clamp) fit function to calculate the relative diffusion time $T_{diff}$ (given with standard error of the fit). Data for three independent experiments are shown.

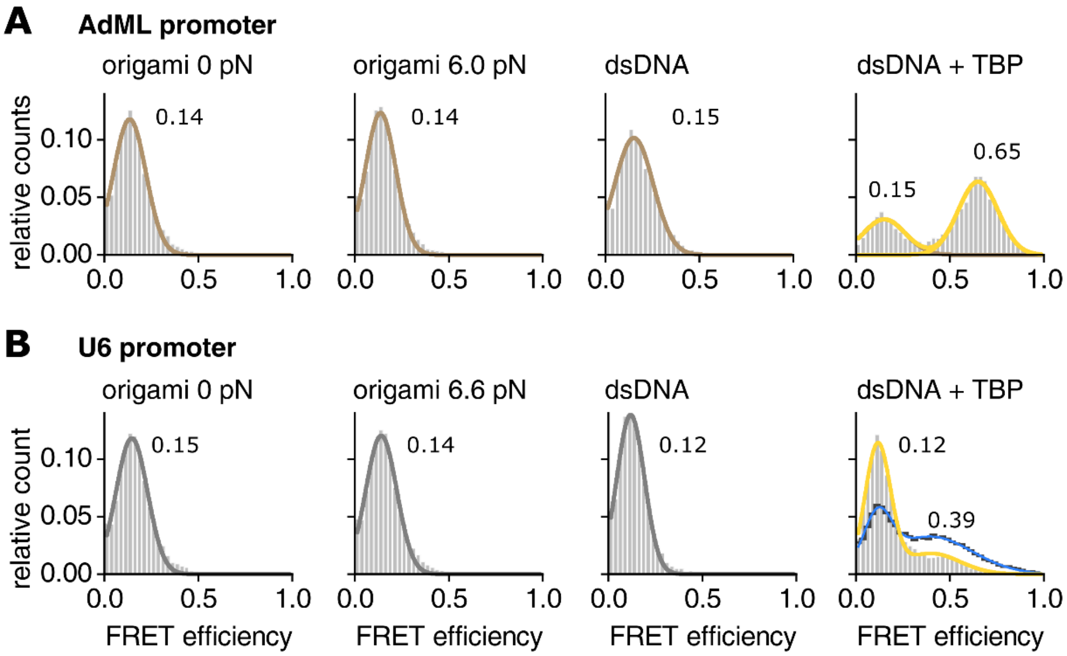


**Supplementary Figure 4: Single-molecule FRET efficiency histograms comparing FRET efficiencies of the DNA origami and linear double stranded DNA oligonucleotides**. The exact same doubly labelled DNA oligonucleotide was used to form a double-stranded promoter DNA as part of the DNA origami or linear double stranded DNA to form the double-stranded **A**) AdMLP or **B**) U6 promoter. The proteins were used in the following concentrations: (A) TBP = 20, (B) TBP = 20 nM (histogram and yellow line for the Gaussian fit), TBP = 100 nM (blue line) – this measurement has been carried out to determine the FRET efficiency of the high FRET state on the U6 promoter (E=0.39) using linear dsDNA. Ds DNA refers to the linear dsDNA without the DNA origami.

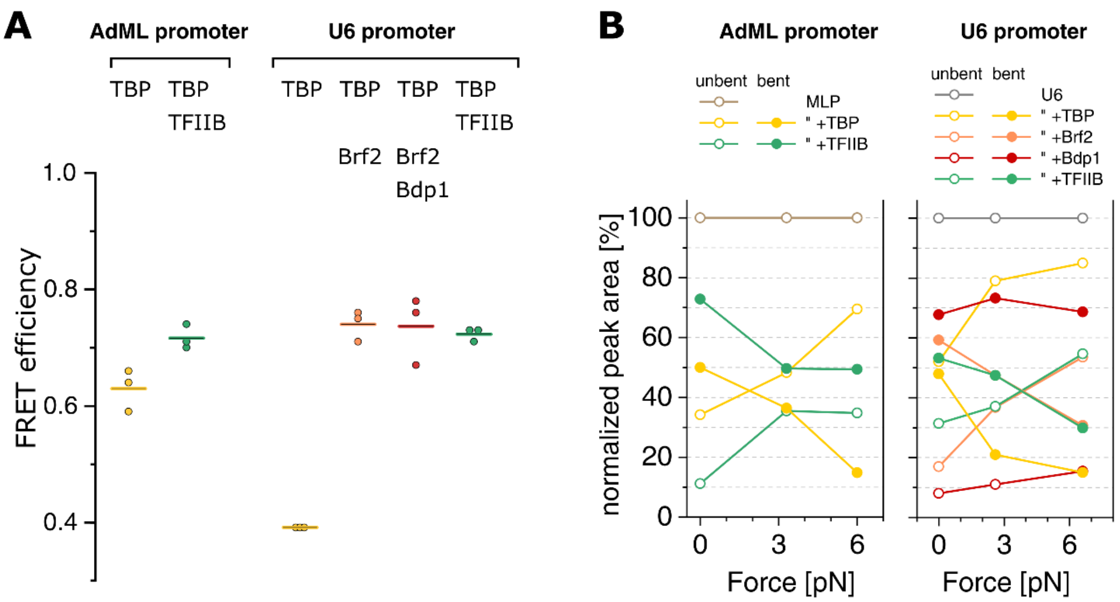


**Supplementary Figure 5: Comparison of FRET efficiencies and peak areas of fitted FRET population for Pol II and III initiation complexes**. **A**) The mean FRET efficiency (straight line) of the high FRET population for all three forces (0, 3.3/2.6, 6.0/6.6 pN, indicated by dots) is summarized. **B**) Changes in fitted peak area of the low FRET (unbent DNA) and high FRET (bent DNA) population under different forces. All areas were normalized to the total fitted area (see **Supplementary Table 4**). The plotted DNA/protein complexes include DNA+TBP (yellow), DNA+TBP+TFIIB (green), DNA+TBP+Brf2 (orange), DNA+TBP+Brf2+Bdp1 (red).

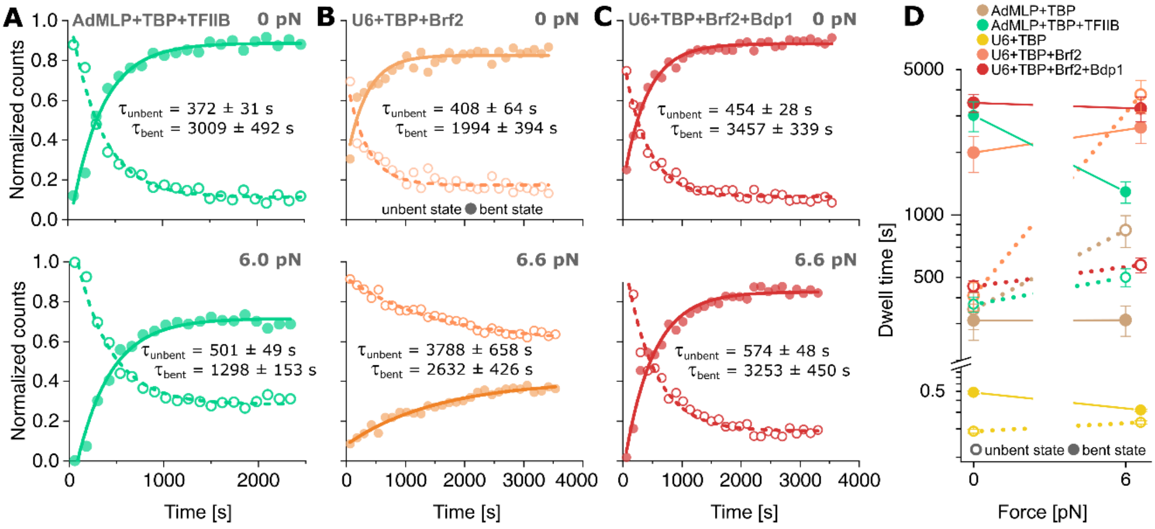


**Supplementary Figure 6: Force dependency of TBP DNA-bending kinetics.**

Relative ratios of low FRET (open circle) to high FRET state (filled circle) from a confocal kinetics experiment with **A)** TBP (20 nM) and TFIIB (200 nM), **B)** TBP and Brf2 (20 nM each) and **C)** TBP, Brf2 and Bdp1 (20 nM each) binding to the AdML promoter at 0 pN and 6.0 pN or the U6 promoter force clamp at 0 pN and 6.0 pN force. Data were fitted with a mono-exponential function. Dwell times were calculated by deconvolution with a perturbation-relaxation model. **D)** Comparison of dwell times in the bent and unbent state for TBP-containing initiation complexes at 0 pN and 6.0 pN (AdMLP) or 6.6 pN (U6) (see also **Figure 4**). Values given as mean ± s.e.m. Connecting lines between data points are meant as visual guides and do not represent interpolations.

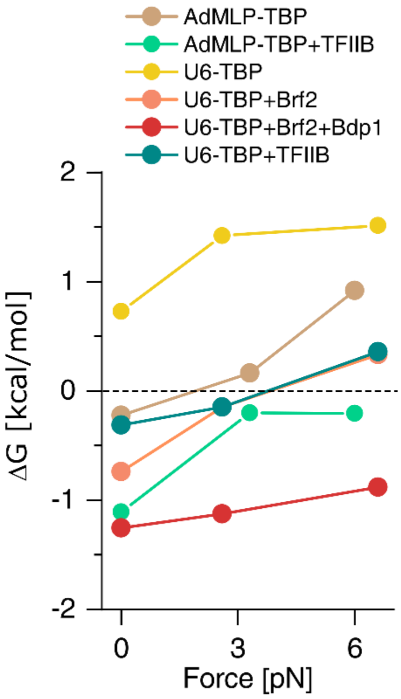


**Supplementary Figure 7: Gibbs free energy of initiation factor complexes under force.**

Complex free energies as a function of force were calculated based on the ratio of the bent state to unbent state (data derived from equilibrium experiments).

**Supplementary Table 3: Fit values for determined for kinetics measurements**. Perturbation-relaxation time course experiments of three technical replicates were combined and the decay of the low FRET population was fitted with a mono-exponential decay function to determine the decay constant and the relative ratio of the low FRET population in dynamic equilibrium y0. Values given are mean ± s.e.m.

| **Complex** | **Force [pN]** | **# Molecules** | **Fit decay constant [s]** | **Fit y0** | **$R^2$ of the fit** |
|---|---|---|---|---|---|
| AdMLP + TBP | 0 | 56851 | 165 ± 29 | 0.53 ± 0.01 | 0.93 |
| “ | 6.0 | 64111 | 228 ± 36 | 0.73 ± 0.01 | 0.87 |
| AdMLP + TBP + TFIIB | 0 | 17570 | 331 ± 24 | 0.11 ± 0.01 | 0.98 |
| “ | 6.0 | 67574 | 361 ± 30 | 0.28 ± 0.01 | 0.98 |
| U6 + TBP + Brf2 | 0 | 23607 | 339 ± 49 | 0.17 ± 0.01 | 0.89 |
| “ | 6.6 | 63193 | 1553 ± 209 | 0.59 ± 0.02 | 0.97 |
| U6 + TBP + Brf2 + Bdp1 | 0 | 44005 | 401 ± 22 | 0.12 ± 0.01 | 0.98 |
| “ | 6.6 | 61715 | 488 ± 36 | 0.15 ± 0.01 | 0.97 |

**Supplementary Table 4: Average FRET efficiencies and peak areas of fitted FRET population for Pol II and III initiation complexes**. FRET efficiency histograms of three technical replicates were fitted with a single (DNA) or triple Gaussian distribution. The average FRET efficiency was determined as the center of each fit population. The area of each peak was normalized to the total fit area. *The value was fixed for fitting. **Peak 2 was included to allow more accurate fitting of the low and high FRET population with a fixed area of 0.004 or *** 0.006.

| | **# Molecules** | **Average FRET efficiency** | | | **Normalized peak area [%]** | | |
|---|---|---|---|---|---|---|---|
| | | Peak 1 (unbent DNA) | Peak 2** | Peak 3 (bent DNA) | Peak 1 (unbent DNA) | Peak 2** | Peak 3 (bent DNA) |
| **AdML promoter** | | | | | | | |
| 0 pN DNA | 27729 | 0.13 | | | 100.0 | | |
| 0 pN TBP | 26187 | 0.12 | 0.37 | 0.64 | 34.3 | 15.7 | 50.0 |
| 0 pN TBP+TFIIB | 36027 | *0.12 | 0.47 | 0.74 | 11.2 | 16.0 | 72.8 |
| 3.3 pN DNA | 40289 | 0.15 | | | 100.0 | | |
| 3.3 pN TBP | 48743 | 0.10 | 0.31 | 0.59 | 48.3 | 15.3 | 36.4 |
| 3.3 pN TBP+TFIIB | 59994 | 0.12 | 0.43 | 0.70 | 35.4 | 14.9 | 49.6 |
| 6 pN DNA | 33930 | 0.14 | | | 100.0 | | |
| 6 pN TBP | 37981 | 0.13 | *0.40 | 0.66 | 69.5 | 15.6 | 14.8 |
| 6 pN TBP+TFIIB | 35307 | 0.13 | 0.40 | 0.71 | 34.8 | 15.8 | 49.4 |
| **U6 snRNA promoter** | | | | | | | |
| 0 pN DNA | 25170 | 0.15 | | | 100.0 | | |
| 0 pN TBP | 31961 | 0.14 | | *0.39 | 0.52 | | 0.48 |
| 0 pNTBP+Brf2 | 33693 | *0.16 | 0.45 | 0.76 | 17.0 | ***23.7 | 59.3 |
| 0 pN_TBP+Brf2+Bdp1 | 30453 | *0.16 | 0.47 | 0.78 | 8.1 | ***24.2 | 67.7 |
| 2.6 pN DNA | 25960 | 0.15 | | | 100.0 | | |
| 2.6 pN TBP | 26820 | 0.14 | | *0.39 | 0.79 | | 0.21 |
| 2.6 pN TBP+Brf2 | 44296 | 0.15 | 0.45 | 0.75 | 36.8 | 15.8 | 47.4 |
| 2.6 N TBP+Brf2+Bdp1 | 25724 | *0.16 | 0.50 | 0.76 | 11.0 | 15.7 | 73.2 |
| 6.6 pN DNA | 49120 | 0.15 | | | 100.0 | | |
| 6.6 pN TBP | 10178 | 0.13 | | *0.39 | 0.85 | | 0.15 |
| 6.6 pN TBP+Brf2 | 45829 | 0.15 | 0.38 | 0.71 | 53.5 | 15.7 | 30.7 |
| 6.6 pN TBP+Brf2+Bdp1 | 48274 | *0.15 | 0.39 | 0.67 | 15.5 | 15.9 | 68.7 |
| 0 pN TBP+TFIIB | 21659 | 0.14 | 0.43 | 0.73 | 31.4 | 15.3 | 53.3 |
| 2.6 pN TBP+TFIIB | 18543 | 0.12 | 0.43 | 0.73 | 37.1 | 15.4 | 47.5 |
| 6.6 pN TBP+TFIIB | 21223 | 0.12 | 0.34 | 0.71 | 54.7 | 15.5 | 29.8 |

### References

1. Nickels, P. C. *et al.* Molecular force spectroscopy with a DNA origami-based nanoscopic force clamp. *Science (New York, N.Y.)* **354,** 305–307; 10.1126/science.aah5974 (2016).
2. Mastronarde, D. N. Automated electron microscope tomography using robust prediction of specimen movements. *Journal of Structural Biology* **152,** 36–51; 10.1016/j.jsb.2005.07.007 (2005).
3. Zivanov, J. *et al.* New tools for automated high-resolution cryo-EM structure determination in RELION-3. *eLife* **7**; 10.7554/eLife.42166 (2018).
